# Force generation by protein-DNA co-condensation

**DOI:** 10.1101/2020.09.17.302299

**Authors:** Thomas Quail, Stefan Golfier, Maria Elsner, Keisuke Ishihara, Vasanthanarayan Murugesan, Roman Renger, Frank Jülicher, Jan Brugués

**Affiliations:** Max Planck Institute of Molecular Cell Biology and Genetics, 01307 Dresden, Germany; Max Planck Institute for the Physics of Complex Systems, 01187 Dresden, Germany; Center for Systems Biology Dresden, 01307 Dresden, Germany; Cluster of Excellence Physics of Life, TU Dresden, 01307 Dresden, Germany

## Abstract

Interactions between liquids and surfaces generate forces^1,2^ that are crucial for many processes in biology, physics, and engineering, including the motion of insects on the surface of water^3^, modulation of the material properties of spider silk^4^, and self-assembly of microstructures^5^. Recent studies have shown that cells assemble biomolecular condensates via phase separation^6^. In the nucleus, these condensates are thought to drive transcription^7^, heterochromatin formation^8^, nucleolus assembly^9^, and DNA repair^10^. Here, we show that the interaction between liquid-like condensates and DNA generates forces that might play a role in bringing distant regulatory elements of DNA together, a key step in transcriptional regulation. We combine quantitative microscopy, *in vitro* reconstitution, optical tweezers, and theory to show that the transcription factor FoxA1 mediates the condensation of a DNA-protein phase via a mesoscopic first-order phase transition. After nucleation, co-condensation forces drive growth of this phase by pulling non-condensed DNA. Altering the tension on the DNA strand enlarges or dissolves the condensates, revealing their mechanosensitive nature. These findings show that DNA condensation mediated by transcription factors could bring distant regions of DNA in close proximity, suggesting that this physical mechanism is a possible general regulatory principle for chromatin organization that may be relevant *in vivo*.

## Main text

Compartmentalization is key to organizing cellular biochemistry. Biomolecular condensate formation underlies the compartmentalization of many cellular functions^6^. Considerable progress has been made towards understanding the biophysical properties of condensates in bulk. However, how these condensates interact with other cellular components such as polymers, membranes, and chromatin remains unclear. Transcriptional hubs are an example of compartments in the nucleus. These hubs involve the coalescence of transcription factors, biochemical regulators of transcription, and DNA^11^. The physical nature of these transcription hubs is under debate, though recent studies have proposed that transcriptional hubs can be understood as examples of biomolecular condensates^12^. In theory, the interactions between transcriptional machinery condensates and the DNA polymer could deform DNA, potentially bridging distal regulatory elements, a critical step in gene regulation. However, we still lack a physical picture of how transcriptional regulators interact with each other and with the surface of the DNA polymer.

To investigate how transcription factors physically organize DNA, we attached linearized λ-phage DNA to a coverslip via biotin-streptavidin linkers (Fig. 1a). We used TIRF microscopy to image the interactions between DNA and Forkhead Box Protein A1 (FoxA1), a pioneer transcription factor that regulates tissue differentiation across a range of organisms^13^ (Fig. 1b). Upon addition of 10 nM FoxA1-mCherry (FoxA1) to the flow chamber in the presence of DNA, FoxA1 formed protein condensates that decorated the strand (Fig. 1c). In the absence of DNA, FoxA1 did not nucleate condensates in solution at concentrations ranging from 10 to 500 nM (Extended Data Fig. 1a). The requirement for DNA in condensate formation at low concentrations suggests that DNA mediates the condensation of a thin layer of FoxA1 on DNA.

**Figure 1:**
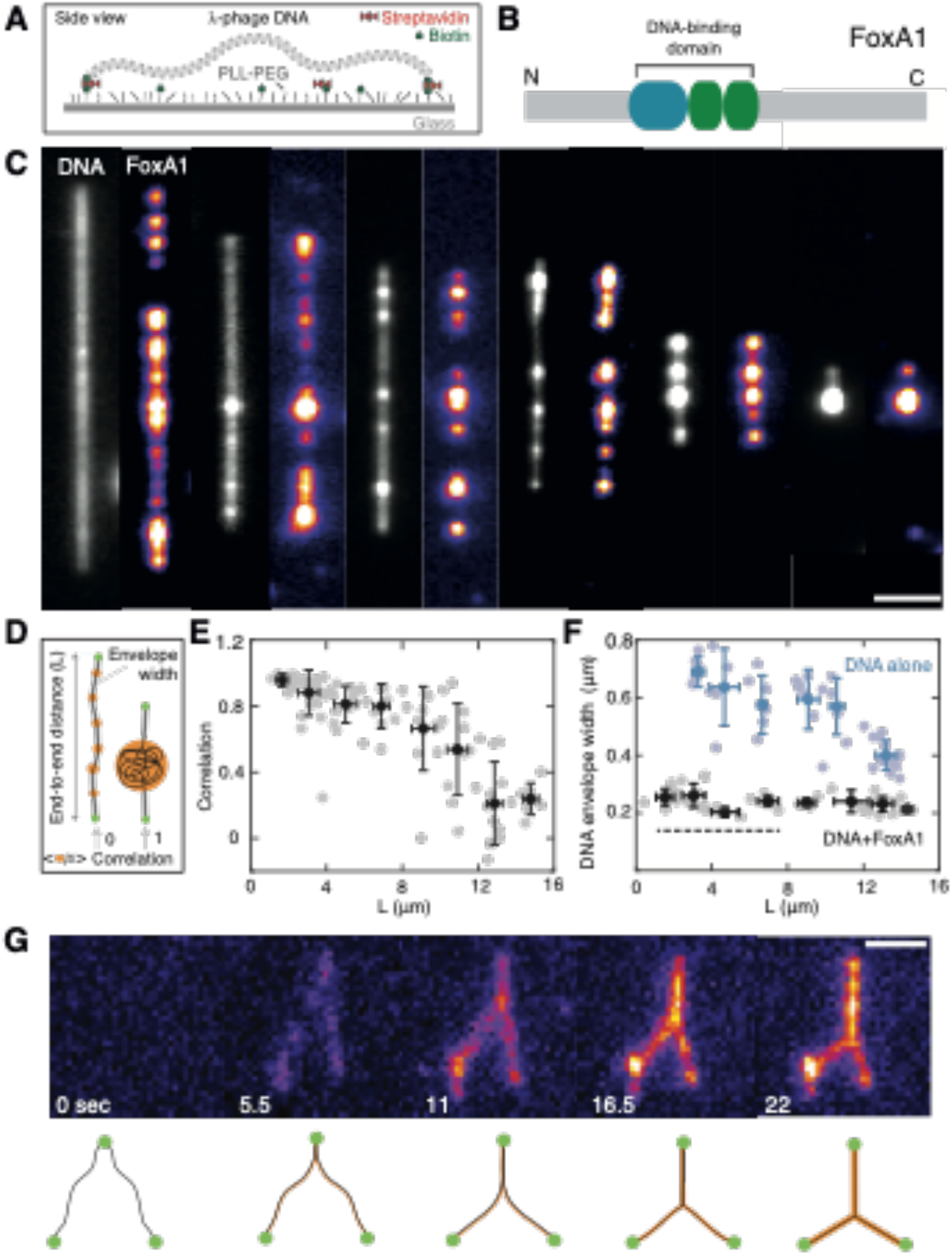
FoxA1 forms DNA-FoxA1 condensates in a tension-dependent manner. (A) Schematic of single λ-phage DNA molecule assay. (B) Structure of FoxA1, consisting of a structured DNA-binding domain flanked by mostly disordered N and C termini. The DNA-binding domain has a sequence-specific binding region (blue) and two non-sequence-specific binding regions (green). (C) The extent of FoxA1-mediated DNA condensation depends on the end-to-end distance of the strand. Representative time-averaged projections of FoxA1 and DNA. Note that the total amount of DNA is the same in each example. The DNA was imaged using 10 nM Sytox Green. Scale bar=2 *μ*m. (D) Schematic displaying three main quantities used to characterize DNA-FoxA1 condensation: the end-to-end distance L; Cross-correlation of DNA and FoxA1 intensities; and DNA envelope width, a measure of transverse DNA fluctuations. (E) Cross-correlation of FoxA1 and DNA signals shows that FoxA1 condenses DNA below a critical end-to-end distance. The gray dots represent individual strands, n=107. The data is binned every 2-*μ*m (black, mean ± SD for both correlations and strand lengths). (F) DNA envelope width measurements (see Methods) reveal that FoxA1-DNA condensation buffers DNA tension (blue and black dots correspond to control and DNA+FoxA1 conditions, n=45 and n=50 respectively). The data is binned every 2- *μ*m (mean ± SD for both the envelope widths and strand lengths). The dashed black line represents the theoretical diffraction limit. (G) Representative images of FoxA1 zipping two independent DNA strands over time. Scale bar=2 *μ*m.

In our assay, DNA molecules displayed a broad distribution of end-to-end distances (L), determined by the DNA-coverslip attachment points (Fig. 1c, d). This end-to-end distance tunes the tension of the DNA^14^. For DNA strands with end-to-end distances greater than approximately 10 μm, FoxA1 generated protein condensates on DNA (Fig. 1c). However, FoxA1 condensation did not influence the DNA molecule (Fig. 1c, leftmost pair of images). Strikingly, for DNA molecules with end-to-end distances below 10 μm, FoxA1 pulled DNA into highly enriched condensates of FoxA1 and DNA (Fig. 1c, Extended Data Fig. 1b-e) with a density of roughly 750 molecules/μm^3^ (see Methods, Extended Data Fig. 2a-d). To quantify FoxA1-mediated DNA condensation, we measured the cross-correlation of FoxA1- DNA intensities as a function of end-to-end distance (see Methods, Fig. 1d,e, Extended Data Fig. 3a). Consistent with the ability of FoxA1 to form FoxA1-DNA condensates at low tensions, the cross-correlation decayed from one to zero with increasing end-to-end distance (Fig. 1e). Thus, FoxA1 mediates the formation of a DNA–protein-rich phase in a tension-dependent manner.

The observation that FoxA1 drives DNA condensation suggests that it can overcome the DNA molecule’s entropic tension set by the end-to-end distance^14^. Incorporating DNA into the condensates increases the tension on the strand, thereby reducing the transverse DNA fluctuations of the non-condensed DNA. To quantify this, we measured the DNA envelope width of the non-condensed DNA fluctuations (see Methods, Extended Data Fig. 3b). In buffer, the DNA envelope width decreased as a function of end-to-end distance, consistent with the corresponding increase of DNA strand tension for increasing end-to-end distances^14^ (Fig. 1f). However, in the presence of FoxA1, the DNA envelope width remained constant for all end-to-end distances as FoxA1 pulled DNA into one or more condensates. The magnitude of the DNA envelope width was lower in the presence of FoxA1 than in buffer conditions for all end-to-end distances (Fig. 1f). Taken together, this suggests that FoxA1-DNA condensates generate forces that can overcome the entropic tension of the non-condensed DNA and buffer its tension.

The observation that FoxA1 can mediate DNA condensation suggests that it could bridge distant DNA strands. To investigate this possibility, we examined DNA molecules that were bound to the same streptavidin molecule at one end (Fig. 1g, Extended Data Fig. 3c). In the absence of FoxA1, these DNA molecules form a v-shaped morphology and fluctuate independently of one another. Upon addition of FoxA1, however, we observed that the two strands zipped together, generating a y-shaped morphology as the condensation of FoxA1 increased over time (Fig. 1g, Extended Data Fig. 3c). Taken together, these data demonstrate that FoxA1 can physically bridge DNA strands in both *cis* and *trans*.

Two mechanisms can be postulated to explain FoxA1-mediated DNA condensation in our experiments: (i) direct cross-linking via the multiple DNA-binding activities of FoxA1^15^ or (ii) weak protein-protein interactions driven by disordered regions of FoxA1. FoxA1 consists of a winged helix-turn-helix DNA-binding domain and two N and C termini domains that are mostly disordered^15^. The DNA-binding domain contains a sequence-specific binding region composed of three alpha helices and a non-sequence-specific binding region composed of two wings. Two point mutations known to affect sequence-specific DNA binding (NH-FoxA1^15^) had virtually no influence on DNA condensation activity (Fig. 2a). Although the presence of two point mutations known to affect non-sequence-specific DNA binding (RR-FoxA1^15^) partially inhibited FoxA1 localization to the strand (Fig. 2b), this mutant still condensed DNA. In this case, condensation occurred on a time scale of minutes rather than seconds (as in WT-FoxA1), which can be explained by the delay in condensing sufficient RR-FoxA1 to the strand. These data suggest that non-sequence-specific binding drives the localization of FoxA1 to DNA but does not mediate DNA condensation through cross-linking. Furthermore, the sequence-specific binding domain of FoxA1 is dispensable for its localization to DNA *in vitro*. To probe whether FoxA1 protein-protein interactions through disordered domains mediate DNA condensation, we truncated both the N and C termini of FoxA1. Although ΔN-FoxA1 retained DNA condensation activity (Fig. 2c), truncating the disordered C terminus of FoxA1 largely inhibited DNA condensation activity (Fig. 2d). Additionally, we found that, at high FoxA1 concentrations in bulk (50 μM), 3% PEG (30K) nucleated highly-enriched spherical FoxA1 condensates (Extended Data Fig. 4a), further suggesting the existence of weak FoxA1-FoxA1 interactions. Thus, non-sequence-specific binding drives FoxA1 localization to DNA, and the disordered C terminus of FoxA1 promotes DNA condensation.

**Figure 2:**
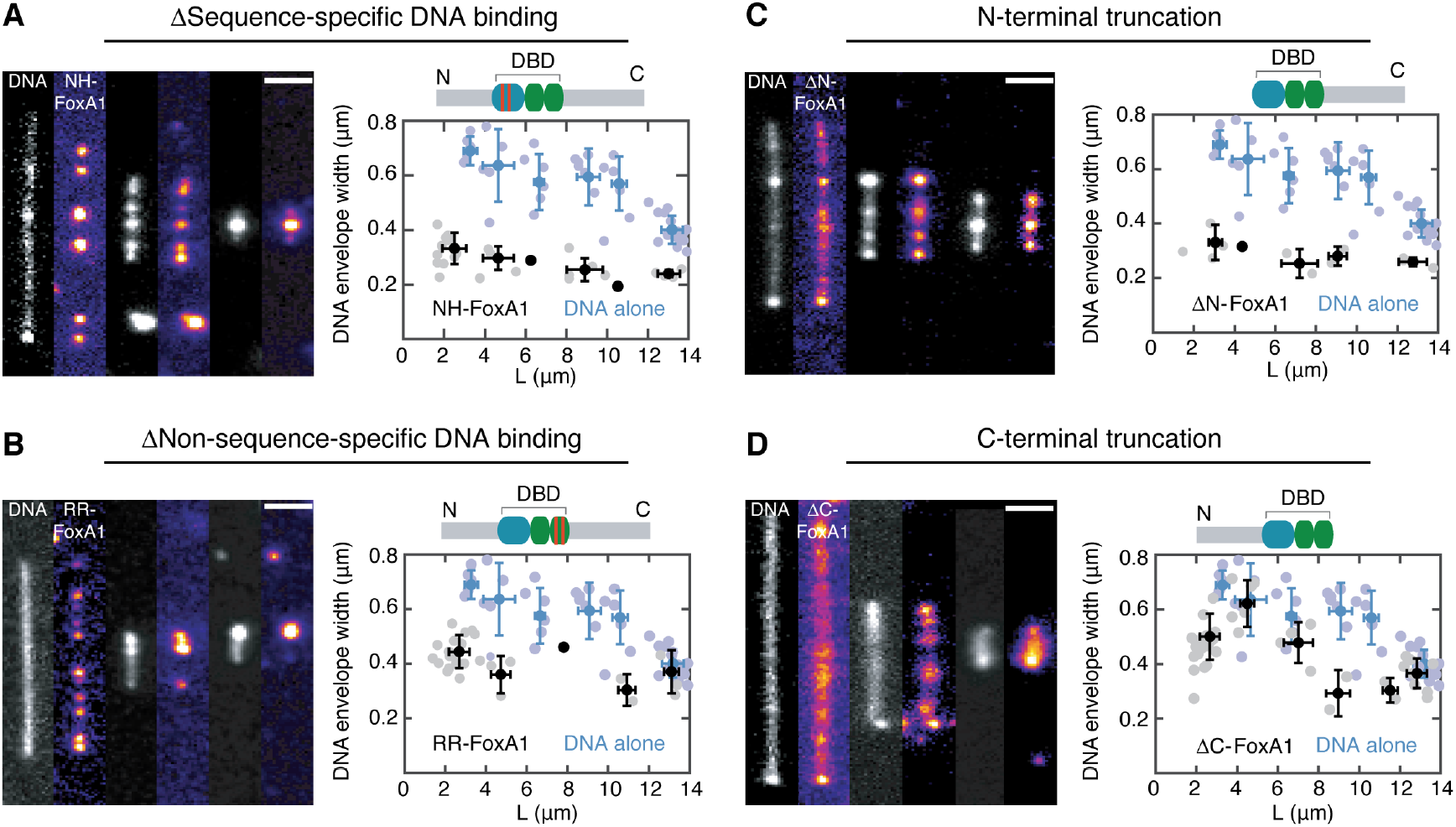
Mutant analysis reveals that the C terminus of FoxA1 drives DNA condensation. Representative images and DNA envelope width measurements for FoxA1 mutants. The data is binned every 2-μm and the mean ± SD (for both the envelope width and the strand length) are shown in black for each mutant and in blue for control (n=45). Scale bars=2 *μ*m. (A) Sequence-specific DNA binding mutant NH-FoxA1 condenses DNA (n=30). (B) Non-sequence-specific DNA-binding mutant RR-FoxA1 condenses DNA (n=28). (C) N-terminal truncation of FoxA1 ΔN-FoxA1 condenses DNA (n=13). (D) C-terminal truncation of FoxA1 ΔC-FoxA1 inhibits DNA condensation (n=44). In all conditions, the protein concentration was 10 nM.

Our results support the hypothesis that FoxA1 condenses onto DNA to generate a DNA–protein-rich condensate via weak protein-protein interactions that exerts a pulling force on the non-condensed strand (see the section Thermodynamic description of DNA-protein condensation in the Supplementary information). To explore the thermodynamics of condensation, we developed a theoretical description based on a semi-flexible polymer partially condensing into a liquid-like condensate. Here, the semi-flexible polymer is DNA and the condensation is mediated by the transcription factor. The free energy of this process contains volume, 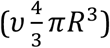, and surface contributions, (*γ*4*πR*^2^), as well as a term representing the free energy of the non-condensed DNA (Fig. 3a), where *ν* is the condensation free energy per volume, *R* is the condensate radius, and *γ* is the surface tension of the condensate. We assume that DNA is fully collapsed inside the condensate and thus its volume is proportional to the condensed DNA contour length, *V* = *α L*_*d*_, where 1/*α* describes the packing density given as DNA length per condensate volume. The free energy of the polymer, 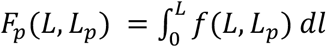, can be obtained from the force-extension curve of the polymer *f*(*L, L*_*p*_), where *L*_*p*_ is the contour length of the non-condensed polymer. Using *L*_*p*_ = *L*_*c*_ − *L*_*d*_ where *L*_*c*_ is the contour length of λ-phage DNA (16.5 μm), the free energy is as follows,

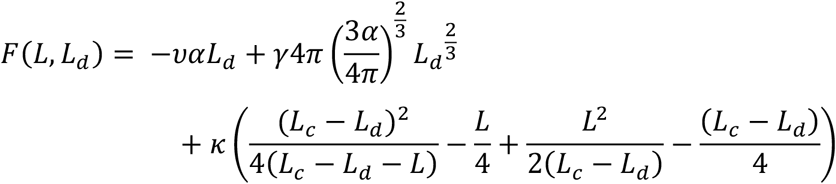

where 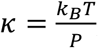, *k*_*B*_ is the Boltzmann constant, T is the temperature, and P is the persistence length of DNA (see the section Thermodynamic description of DNA- protein condensation in the Supplementary information). For fixed *L*, the minimum of *F*(*L, L*_*d*_) determines the preferred size of the condensate. This free energy predicts upon variation of *L* a stochastic first-order phase transition for the formation of DNA–protein condensates (Fig. 3b). The distribution of condensate sizes is then given by 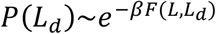, for fixed *L* (Fig. 3c). This accounts for a sharp transition of DNA condensation controlled by the end-to-end distance and thus the tension of the DNA molecule. The first-order nature of this behavior implies regimes of hysteresis and bistability. Our theory also predicts that the condensation forces exerted on the non-condensed DNA are kept roughly constant.

**Figure 3:**
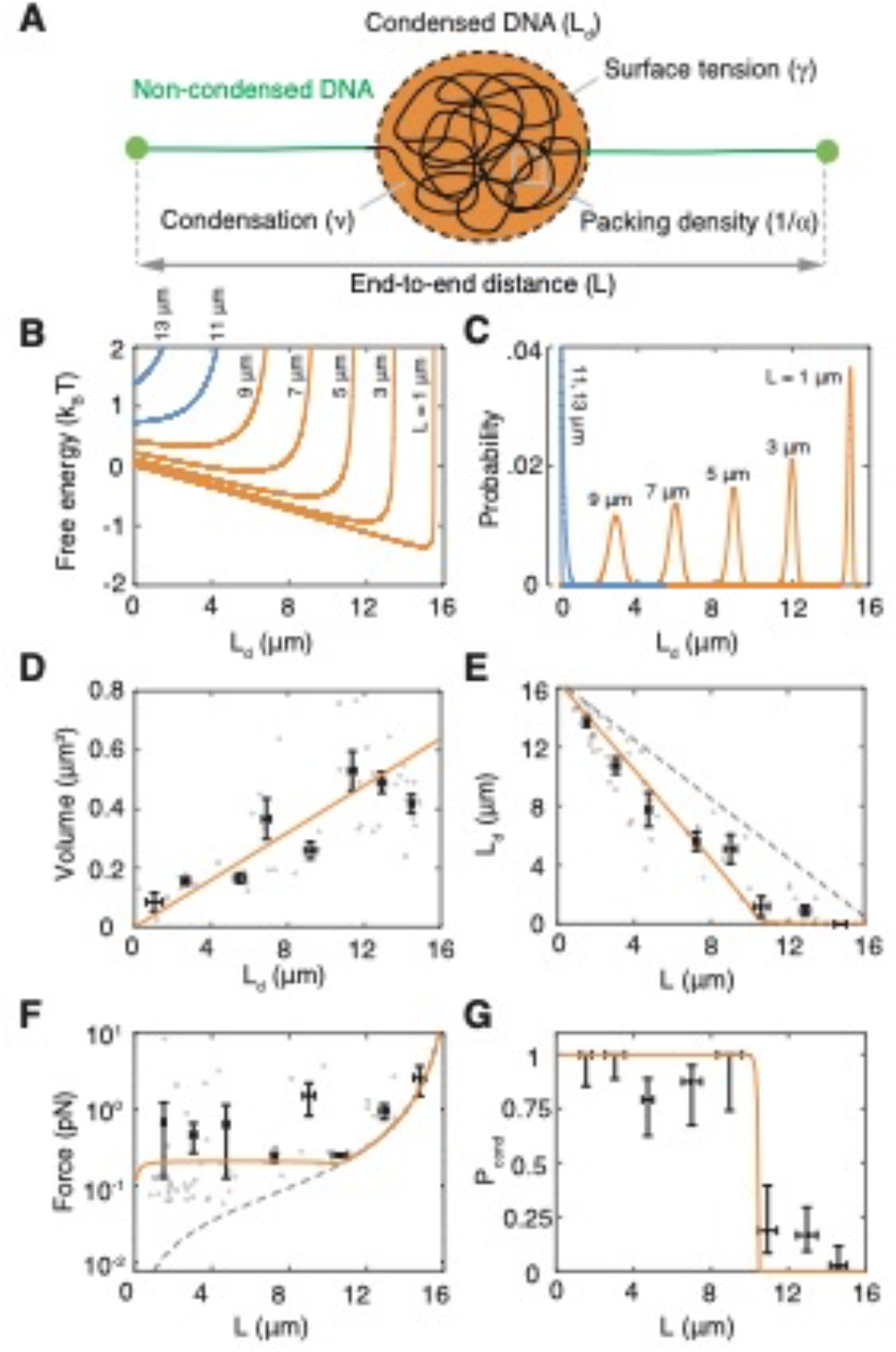
Thermodynamic description of a liquid phase condensing onto a semi-flexible polymer explains FoxA1-mediated DNA condensation. (A) Schematic representing DNA-FoxA1 condensation (orange). DNA can be in a condensed state (black) or a non-condensed state (green). DNA condensation depends on the condensate surface tension (*γ*), condensation free energy per volume (*ν*), and DNA packing efficiency (α). (B) Free energy profiles as a function of condensed DNA (L_d_) for different L reveal a first-order phase transition for DNA–protein condensation (orange and blue correspond to favorable and unfavorable condensation, respectively). (C) Boltzmann distributions corresponding to the free energy profiles in (B). (D) Condensate volume linearly increases with L_d_. The orange curve represents a linear fit to individual strands (n=47). For (D), (E), and (F), individual strands are represented as gray dots and binned mean±SEM is in black. (E) Amount of condensed DNA as a function of L (n=63) reveals sharp transition. Orange curve represents optimal theoretical fit. The gray dashed-line corresponds to the limit of maximum condensation where L_d_ is equal to the contour length of DNA (16.5 μm) minus L. (F) Condensation forces that DNA-protein condensates exert on non-condensed DNA are buffered (n=62). Orange curve is the theoretical prediction. The gray dashed line represents the force when L_d_=0. (G) Probability to nucleate a DNA-FoxA1 condensate (P_cond_) reveals a sharp transition at a critical end-to-end distance. P_cond_ is computed from binned local correlation data (n=181 condensates). The end-to- end distance error bars are the SD and the P_cond_ error bars are the 95% confidence intervals from a Beta distribution.

To test this theory, we first measured DNA condensate volumes and found that they increase linearly with the length of condensed DNA (*L*_*d*_), with *α*=0.04 ± 0.01 μm^2^ (Fig. 3d, Extended Data Fig. 4d, Methods). This confirms that DNA is in a collapsed conformation inside the condensates. Next, we simultaneously fit the predictions to the average amount of DNA contained in the condensates (*L*_*d*_), and the probability of nucleating a DNA condensate (*P*_*cond*_) as a function of end-to-end distance (see Methods). We calculated *L*_*d*_ (Fig. 3e, Extended Data Fig. 4e, Extended Data Fig. 5) and *P*_*cond*_ (Fig. 3g, Extended Data Fig. 4f) using the Boltzmann probability distributions (Fig. 3c) from the free energy. Our fits agree quantitatively with the data and show that *L*_*d*_ decreases with *L* until a critical end-to-end distance beyond which DNA condensates do not form. Below this critical length, we observed that the force exerted by the condensate is buffered at 0.21 pN (0.18 – 0.30 pN CI), consistent with the theory (Fig. 3f). To complement our force measurements, we performed optical tweezer measurements of FoxA1- mediated DNA condensation. Incubating a single λ-phage DNA molecule at either *L* = 6 or 8 μm in the presence of 150 nM FoxA1 generated forces on the order of 0.4-0.6 pN, consistent with the force measurements using fluorescence microscopy (Methods, Extended Data Figs. 6,7). Finally, *P*_*cond*_ exhibits a sharp transition at *L* = 10.5 μm (9.4 – 10.9 μm CI), in agreement with a stochastic first-order phase transition (Fig. 3g). We also observed a sudden force jump during the onset of condensate formation (as measured by the individual temporal force trajectories in the optical tweezer experiments), consistent with a first order phase transition (Extended Data Figs. 6c,7). Close to the transition point FoxA1- mediated DNA condensation displayed bistability. This bistability was observed in strands that contained multiple FoxA1 condensates, but where only some of them condensed DNA (Extended Data Fig. 8a). Our fits allowed us to extract the physical parameters associated with condensate formation, namely the condensation free energy per volume *ν* = 2.6 pN/μm^2^ (2.3 – 5.2 pN/μm^2^ CI) and the surface tension *γ*=0.04 pN/μm (0.04 – 0.28 pN/μm CI), see Methods section. These parameters are consistent with previous measurements for *in vitro* and *in vivo* condensates^16,17^.

Our theory and experiments show that two key parameters govern DNA–protein co-condensation, namely the condensation free energy per volume (*ν*) and the surface tension (*γ*). We reasoned that different DNA-binding proteins may exhibit a range of behaviors depending on these parameters. First, we investigated the sequence-specific DNA-binding region mutant (NH-FoxA1), which also condensed DNA but to a lesser extent (Fig. 2a). Quantitatively, we found that the surface tension of condensates formed with this mutant was roughly unchanged compared to WT-FoxA1, *γ*=0.065 pN/μm (0.05 – 0.07 pN/μm CI), but the free energy per volume of condensation was reduced consistent with reduced DNA binding, *ν*=1.05 pN/μm^2^ (0.9 – 1.1 pN/μm^2^ CI), Extended Data Fig. 9, Fig. 4a. This was also reflected in a decrease in the extent of DNA packing with *α* = 0.09 ± 0.02 μm^2^ (Extended Data Fig. 9a). We also observed that NH-FoxA1-mediated condensates generated a force of 0.17 pN (0.16 – 0.19 pN, CI), lower than that for WT-FoxA1. In addition, NH-FoxA1 displayed bistable DNA–protein condensation activity in the neighborhood of the transition point (Extended Data Fig. 8b). Next, we examined the interactions of a different transcription factor Tata-Box-binding protein (TBP) with DNA. We found that TBP also formed small condensates on DNA, but did not condense DNA even at the lowest imposed DNA tensions (Fig. 4b). Instead, TBP performed a diffusive motion along the DNA strand (Extended Data Fig. 10c), suggesting that DNA-protein condensation is not thermodynamically favored. Another transcription factor, Gal4-VP16, formed condensates on DNA and condensed DNA in a tension-dependent manner consistent with FoxA1 (Extended Data Fig. 10e). Lastly, we analyzed somatic linker histone H1, a protein that is structurally similar to FoxA1. However, in contrast to FoxA1, one of the known functions of H1 is to compact chromatin^18^, so we expected H1 to strongly condense DNA. Consistent with this, we found that H1 displayed a stronger DNA condensation activity compared to FoxA1, condensing DNA for all measured end-to-end distances (Fig. 4c). Interestingly, the *Xenopus* embryonic linker histone B4 condensed DNA in a tension-dependent manner but not to the same extent as H1 (Extended Data Fig. 10f). Thus, we propose that the competition between condensation free energy per volume of the DNA–protein phase and surface tension regulate a spectrum of DNA condensation activities, which may be tuned by the structure of transcription factors.

**Figure 4:**
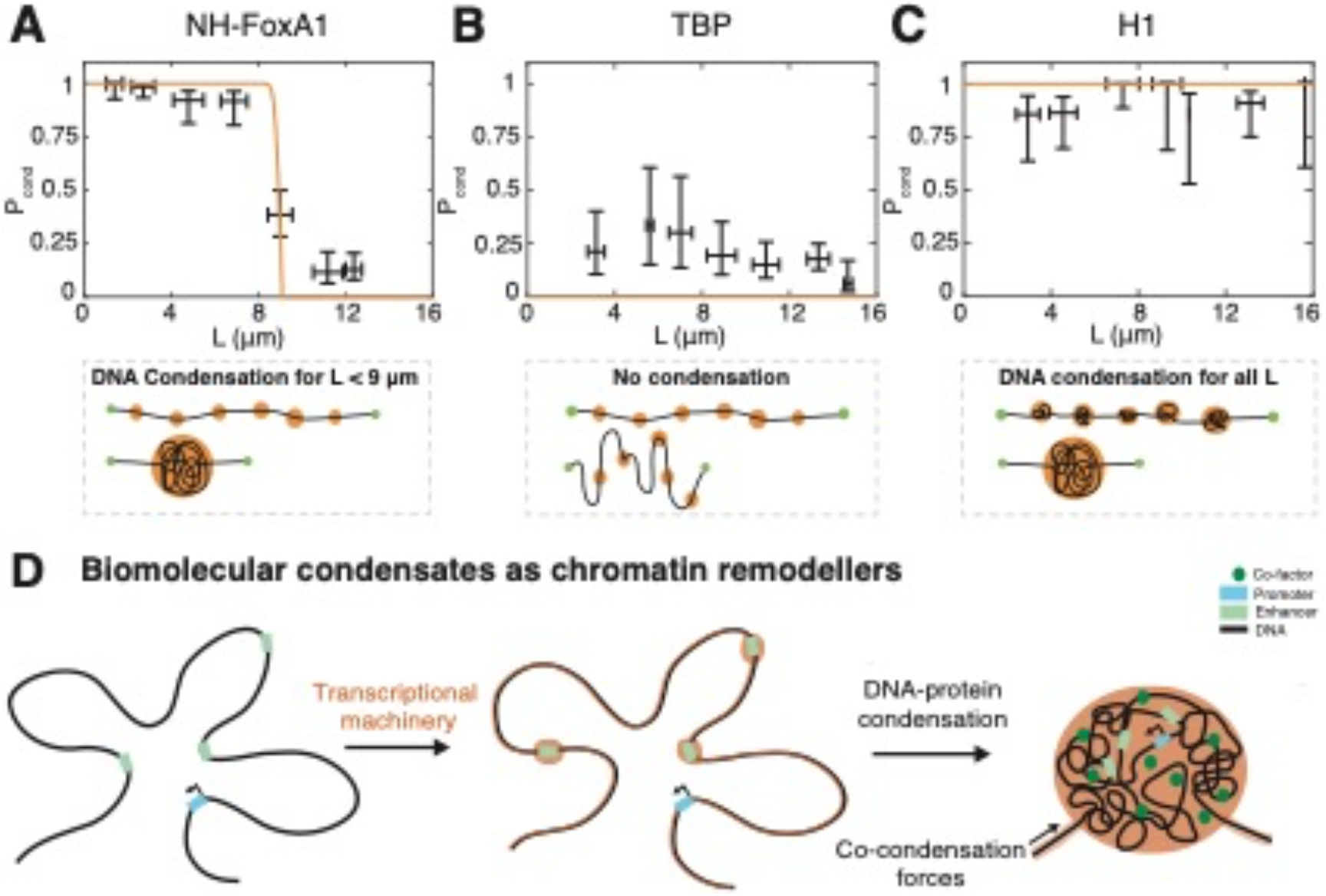
Universality of protein-DNA co-condensation. Probability to form a protein-DNA co-condensate for NH-FoxA1 (A), Tata-box-binding protein (B), and Somatic linker histone H1 (C). P_cond_ is computed from local correlation data with n=361 condensates for NH-FoxA1 (A), n=247 condensates for Tata-box-binding protein (B), and n=101 for H1 (C). Scale bar=2 *μ*m. The error bars for the end-to-end distance are SD and the P_cond_ error bars are the 95% confidence intervals from a Beta distribution. We found that NH-FoxA1 condensed DNA less strongly than WT-FoxA1, TBP could not condense DNA for any end-to-end distance, and H1 condensed DNA for all measured end-to-end distances. (D) Biomolecular condensates generate condensation forces that could serve to recruit transcriptional regulators, and potentially remodel chromatin at physiologically relevant force scales in order to properly regulate transcription. See Figure 2 in the Supplementary Information for representative protein-DNA images of NH- FoxA1, TBP, and H1.

Here, we show that FoxA1 can condense DNA under tension to form a DNA– protein-rich phase that nucleates through a force-dependent first-order transition for forces below a critical value. This critical force, which is on the order of 0.2-0.6 pN for FoxA1, is set by co-condensation forces that the DNA–protein phase exerts on the non-condensed DNA. These forces are similar in magnitude to those recently measured for DNA loop extrusion on the order of 0.2-1 pN^19,20^ and those estimated in intact nuclei from nuclear condensate fusion^21^. Thus, we speculate that these weak forces we find *in vitro* may be of relevance to the mechanics of chromatin organization, though future studies are necessary to show this. Taken together, our work suggests that co-condensation forces may act as an additional mechanism to remodel chromatin in addition to molecular motors that extrude loops and complexes that remove or displace nucleosomes (Fig. 4d).

Transcription-factor-mediated DNA–protein condensation represents a possible mechanism by which transcription factors coordinate enhancer-promoter contacts in transcriptional hubs^12^. In this context, DNA–protein condensates could act as scaffolds, pulling co-factors into the droplet (Fig. 4d). Our theoretical description reveals that these DNA–protein condensates are formed via a first-order phase transition, suggesting that they can be assembled and disassembled rapidly by changing mechanical conditions. Near the transition point, assembly and disassembly of these *in vitro* DNA–protein condensates becomes highly stochastic, reminiscent of the rapid dynamics associated with the initiation and cessation of transcriptional bursts observed in vivo^22^.

We have demonstrated that protein-DNA co-condensation is associated with a difference in chemical potential between the condensed and non-condensed DNA. This difference in chemical potential is transduced by the condensate to perform mechanical work on the non-condensed DNA strand. Capillary forces represent another example of forces that involve liquid-surface interactions^1,2,23^. With both co-condensation and capillary forces, attractive interactions give rise to the transduction of free energy into work. Such forces may also be relevant beyond chromatin in other biological contexts, including membranes and the cytoskeleton.

DNA–protein co-condensation not only provides mechanisms to facilitate enhancer–promoter contacts, but could also play a more general role in DNA compaction and maintenance of bulk chromatin rigidity in processes such as mitotic chromatid compaction^24^, and the formation of chromatin compartments^8,25,26^. Owing to the tension-dependent nature of DNA–protein co-condensation, our work suggests that these forces could play a key and, as yet, underappreciated role in genome organization and transcriptional initiation. It is appealing to imagine that transcriptional outputs not only respond to concentrations of transcription factors in the nucleus, but also to mechanical cues from chromatin.

## Acknowledgements

We thank Pavel Tomancak, Anthony Hyman, Stephan Grill, Iain Patten, Martin Loose, David Oriola, Benjamin Dalton, Patrick McCall, and Claudia Meyer for helpful feedback and stimulating discussions. We would like to thank Aliona Bogdanova for discussions and help with cloning as well as both the Protein Expression and Purification Facility and the Light Microscopy Facility at the Max Planck Institute of Molecular Cell Biology and Genetics. We would like to acknowledge and thank Nadine Vastenhouw for discussions and the construct containing the Tata-box-binding protein template, and Christoph Zechner for help with statistical analyses. Lastly, we would like to thank Kenneth Zaret for sending constructs, information on mutant FoxA1 proteins, and advice on FoxA1 protein purification. This work was supported by an EMBO long-term fellowship (ALTF-1456-2015) (TQ), DFG project BR 5411/1-1 (JB,VN), and a Volkswagen “Life” grant number 96827 (JB,TQ).

## Author contributions

T.Q. and J.B. conceived the project. T.Q. and S.G. performed imaging experiments. S.G. established the single-strand DNA assay. T.Q. purified proteins, made constructs, and performed data analysis. T.Q., J.B., and F.J. performed theoretical calculations. M.E. made the TBP and Gal4-VP16 constructs and purified the proteins. V.M. purified B4. T.Q. and R.R. performed optical tweezer measurements. S.G. and R.R. performed data analysis and contributed to methods writing. K.I. made the initial FoxA1 construct and provided key biochemical support. J.B. and F.J. supervised the work. T.Q., J.B., and F.J. wrote the manuscript and all authors contributed ideas and reviewed the manuscript.

## Methods

### Cloning and protein purification

FoxA1-mCherry was introduced into a bacterial expression vector with an N- terminal His6 tag using Gateway cloning. Unlabeled FoxA1 was cloned and purified the same way. This vector was transformed into T7 express cells (enhanced BL21 derivative, NEB C2566I), grown to OD∼0.4-0.8, whereupon we added 1 mM IPTG and expressed His6-FoxA1-mCherry for 3-4 hours at 37°C. We thawed frozen pellets in binding buffer (1xBB) that contained 20 mM Tris-HCl (pH=7.9), 500 mM NaCl, 20 mM Imidazole, 1 mM MgCl2, supplemented with protease inhibitors and Benzonase. The redissolved pellets were lysed and clarified via centrifugation. Discarding the supernatant, we resuspended the pellets in 1xBB + 6 M Urea, spun, collected the supernatant and poured it over an IMAC column, eluting the protein with 1xBB+6 M Urea+250 mM Imidazole. We dialyzed overnight into storage buffer (1xSB), 20 mM HEPES (pH=6.5), 100 mM KCl, 1 mM MgCl2, 3 mM DTT, and 5 M Urea. Multiple dialysis rounds reduced the concentration of urea. Finally, the protein was dialyzed into 1xSB+2 M Urea, spun-concentrated to 4-5 mg/ml (∼50 μM), and then snap-frozen nitrogen and stored at -80°C. NH-FoxA1-mCherry and RR-FoxA1-mCherry were obtained following^15^ using the Q5 Site-Directed Mutagenesis Kit. The truncation constructs were generated using restriction digestion-ligation approaches coupled with PCR. We used Alexa-488-labeled somatic linker histone H1 purified from calf thymus (H-13188, ThermoFisher). To purify mCherry-B4, the gene (Genscript) was cloned into a bacterial expression vector with N-terminal His6 and mCherry tags, transformed into T7 express cells, grown to OD∼0.7 and supplemented with 0.8 mM IPTG and expressed at 37°C for four hours. Resuspending the pellets in lysis buffer, 1xPBS with 500 mM NaCl, 1 mM DTT plus protease inhibitors and Benzonase, we then lysed the cells, collected the supernatant, ran the supernatant over an IMAC column, and eluted the protein with lysis buffer+250 mM Imidazole. The protein was dialyzed into 1xPBS+500 mM NaCl overnight, spun-concentrated, snap-frozen, and stored at -80 °C. We purified labeled versions of Tata-box binding protein and Gal4-VP16 using similar purification strategies. Both vectors—His6-MBP-eGFP-zTBP and His6-Gal4-GFP- VP16—were transformed into T7 express cells, grown to OD∼0.6, whereupon we added 0.2 mM IPTG and expressed overnight at 18 °C. We lysed the cells into buffer containing 50 mM Tris-HCl (pH=8.0), 1 M NaCl, 10% glycerol, 1 mM DTT, 1 mM MgCl2 supplemented with protease inhibitors. For subsequent steps, 10 μM ZnSO4 was added to buffers for the Gal4-VP16 purification. After lysis, we added NP40 to 0.1% and clarified via centrifugation. We performed a polyethylenimine precipitation to precipitate DNA and then an ammonium sulfate precipitation to recover the protein, resuspending the precipitated proteins in buffer containing 50 mM Tris-HCl (pH=8.0), 1 M NaCl, 10% glycerol, 1 mM DTT, 0.1% NP40, and 20 mM imidazole and clarified the soluble fraction via centrifugation. We poured the lysate over an IMAC column and eluted the protein using 2xPBS, 250 mM imidazole, 10% glycerol, and 1 mM DTT. We pooled protein fractions and dialyzed TBP overnight into 20 mM HEPES pH=7.7, 150 mM KCl, 10% glycerol, and 1 mM DTT and Gal4-VP16 into 20 mM Hepes (pH=7.7), 100 mM KCl, 50 mM Sucrose, 0.1 mM CaCl2, 1 mM MgCl2, 1 mM DTT, and 10 μM ZnSO4. We then spun-concentrated the proteins, snap-froze using liquid nitrogen, and stored at -80°C.

### DNA functionalization, cover slip PEGylation, and DNA micro-channel preparation

To biotinylate DNA purified from λ-phage (λ-phage DNA), we followed the protocol given in^19^. Each end of the biotinylated λ-phage DNA had two biotin molecules. To PEGylate the cover slips and prepare the DNA microchannels we followed the protocol given in^19^.

### DNA and protein imaging

We fluorescently stained immobilized DNA strands with 10 nM Sytox Green in Cirillo buffer (20 mM HEPES, pH=7.8, 50 mM KCl, 2 or 3 mM DTT, 5% glycerol, 100 μg/ml BSA). For experiments with H1 and TBP, we imaged DNA using 25 nM Sytox Orange. We used protein concentrations of 10 nM. We used a Nikon Eclipse microscope with a Nikon 100x/NA 1.49 oil SR Apo TIRF and an Andor iXon3 EMCCD camera using a frame-rate of 100 – 300ms. A highly inclined and laminated optical sheet (HILO) was established using a Nikon Ti-TIRF-E unit mounted onto the microscope stand.

### Optical tweezer measurements

We performed optical tweezer experiments using a C-Trap G2 system (Lumicks) in a microfluidics flowcell (Lumicks), providing separate laminar flow channels. For each experiment, we trapped two streptavidin-coated polystyrene beads (Spherotec SVP-40-5). Once trapped, we moved these beads to a channel containing biotinylated λ-phage DNA (Lumicks) at a concentration of 0.5 μg/ml, whereupon we used an automated “tether-finder” routine to capture a single molecule between the two beads. Once a single λ-phage DNA molecule was attached to the two beads, we moved the trapped beads to a buffer-only channel (containing Cirillo buffer: 20 mM HEPES, pH=7.8, 50 mM KCl, 3 mM DTT, 5% glycerol, 100 μg/ml BSA). In the buffer-only channel, we fixed the molecule’s end-to-end distance at either L=6 or 8 μm. We then moved the tethered DNA to a channel containing 150 nM FoxA1 in Cirillo buffer or another buffer-only channel (as a control) and tracked the force and imaged the FoxA1-mCherry fluorescence for 100 seconds.

### Bulk phase separation assays

We performed bulk phase separation assays with FoxA1-mCherry, NH-FoxA1- mCherry, and somatic linker histone H1. The storage buffer for FoxA1 and NH- FoxA1 was 20 mM HEPES (pH=6.5), 100 mM KCl, 1 mM MgCl2, 3 mM DTT, and 2 M Urea. The storage buffer for H1 was 1xPBS. For FoxA1, we combined 6 μl of FoxA1 (at 50 μM) and 1 μl of 20% 30K poly-ethylene glycol (PEG). For NH-FoxA1, we combined 9 μl and 1 μl of 20% 30K PEG. For H1, we combined 9 μl H1 and 1 μl 100 μM 32-base pair ssDNA. We prepared flow channels with double-sided tape on the cover slide and attached a PEGylated cover slip to the tape. We imaged the condensates using spinning disk microscopy and a 60x objective.

### FoxA1 molecule number estimation

To estimate the number of FoxA1-mCherry molecules per condensate, we quantified the intensity of single FoxA1-mCherry molecules bound non-specifically to the slide. Around each segmented spot of DNA-independent FoxA1 intensity, we cropped an area of 10×10 pixels, performed a background subtraction and summed the remaining intensity in the cropped area. To determine the contribution of the background, the same method was applied to 10×10 pixel areas void of FoxA1 signal intensity. The resulting distribution of these integrated signal intensities reveals consecutive peaks that are evenly spaced by an average intensity of about 400 a.u., allowing us to calculate the number of molecules. This approach should interpreted as a lower bound estimate of the number of FoxA1-mCherry molecules per condensate, as it neglects effects such as fluorescent quenching^27^.

### Hydrodynamic stretching of DNA

DNA molecules bound at only one end to the slide were hydrodynamically stretched using a constant flow rate of 100 μl/min of 0.5 nM FoxA1-mCherry in Cirillo buffer with 10 nM Sytox Orange. The flow rate was sustained for tens of seconds using a programmable syringe pump (Pro Sense B.V., NE-501).

### Strand length calculation

To calculate the end-to-end distance, we generated time-averaged projections of FoxA1 and DNA and integrated these projections along the strand’s orthogonal axis. To find the profile’s “left” edge, we computed the gradient of the signal and determined the position where the gradient went through a threshold (defined as 0.2). We then took all the points from the start of the signal to this position, performed a background subtraction, and fit an exponential to these points. To ensure that we included the entire DNA signal, we defined the fitted threshold for both the left and the right edges as three-quarters of the value of the fitted exponential value at the point when the gradient had gone through the intensity threshold. Using this fitted threshold, we computed the position values for the left and the right sides, and computed the end-to-end distance as the difference between these two positions.

### Global cross-correlation analysis

We generated time-averaged projections from movies of both FoxA1 and DNA, and then summed the intensities in the orthogonal axis to the strand, generating line profiles. We then calculated the strand length and cropped both the FoxA1 and DNA line profiles from the edges of the strand. We then subtracted the mean value from these cropped line profiles, normalized the amplitudes of the signals by their Euclidean distances, and computed the zero-lag cross-correlation coefficient of the normalized signals, which we defined as “Correlation”: 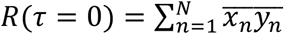,, where *τ* is the number of lags, *N* is the number of points in the normalized FoxA1 and DNA signals, 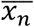, is the *n*th entry of the normalized FoxA1 signal, and 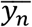, is the *n*th entry of the normalized FoxA1 signal. In general, Correlation values range from -1 to 1, but in our experimental data the values range from roughly 0 to 1, where 1 represents the formation of DNA-FoxA1 condensates and 0 represents the formation of only FoxA1 condensates (no DNA condensation).

### DNA envelope width calculation

To compute the DNA envelope width, we first generated time-averaged projections from movies of FoxA1 and DNA. We then selected segments of the strand that did not contain FoxA1—regions of non-condensed DNA. Using these segments, we extracted a line profile of the DNA signal orthogonal to the strand that gave the maximum width. We then subtracted off the background of the DNA profile, normalized the signal’s amplitude using the Euclidean distance, and fit a Gaussian. We defined the DNA envelope width as 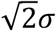,, which represents the square root of two times the standard deviation of the fitted Gaussian. The theoretical diffraction limit is calculated using the Rayleigh criterion, a measure of the minimal resolvable distance between two point sources in close proximity for a given set of imaging conditions: 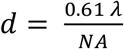,, where *λ* represents the imaging wavelength and NA is the numerical aperture. For our imaging setup, *d* = 0.2 μm, which is approximately 2*σ* of the fluorescent source from the DNA. As the DNA envelope width is defined as 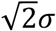,, our “diffraction limit” as given by the dashed line in Fig. 1f is given as 0.14 μm.

### Condensate volume analysis

To calculate condensate volumes, we generated time-averaged DNA-FoxA1 projections and then localized the peaks of the DNA condensates. Using the peak locations, we extracted background-subtracted one-dimensional profiles of the DNA condensates in the orthogonal axis to the strand—these profiles went through the peak location. We fit Gaussians to these profiles without normalizing the amplitude. To define the radii of the condensates, we computed the gradient of the fitted Gaussians and defined the condensate “edges” as when the absolute value of the gradient of the Gaussian function gradient went through a threshold value (defined as one, and determined by comparing with fluorescence). Assuming condensates are spherical, we computed the condensate volume as 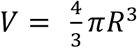, where *R* is the condensate’s radius. To compute a condensate volume for strands with multiple condensates, we simply added up the volumes for each condensate.

### Condensed DNA amount analysis

To compute the amount of condensed DNA, *L*_*d*_, we generated time-averaged projections of DNA and FoxA1 signals, integrating the DNA signal in the orthogonal direction to the strand. We then defined condensed vs non-condensed DNA with Thresholddrop: the median value of the profile plus a tolerance. Intensity values below Thresholddrop were defined as pixels of non-condensed DNA, and intensity values above Thresholddrop were defined as pixels of condensed DNA. This assumption was also consistent with the measured FoxA1 signal, where FoxA1 signals clearly localized to regions of condensed DNA, as defined by the Thresholddrop. The tolerance value was used to suppress artefactual fluctuations of the non-condensed DNA signal in the neighborhood of the median. To optimize the tolerance value, we assume that *L*_*d*_ as a function of *L* is linear for lower values of *L* (<5 *μ*m) with a y-intercept equal to the contour length of the DNA molecule (16.5 *μ*m), as this is consistent with our theoretical description. We plotted the y-intercepts of the linear fits as a function of tolerance and found that tolerance=500 gives a y-intercept equal to 16.5 and generates DNA-FoxA1 condensates up to 10 *μ*m consistent with our data and analysis (Extended Data Fig. 6). To calculate the DNA length contained within the droplet, we integrated the intensities from pixels above Thresholddrop, divided this value by the sum of the total intensity of the profile, and then multiplied this ratio by the contour length of λ-phage DNA, 16.5 *μ*m. The non-condensed DNA length was calculated as simply the contour length minus L_d_. We used the same tolerance = 500 for the NH-FoxA1 mutant analysis.

### Force analysis

To calculate the force that the condensate exerts on the non-condensed DNA, we used the worm-like chain model, which relates λ-phage DNA’s extension and force. Upon addition of FoxA1, the amount of non-condensed DNA reduces, and the extension changes as follows, 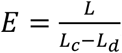,, where *L*_*d*_ is the amount of condensed DNA, *L* is the end-to-end distance, and *L*_*c*_ is the total contour length of the molecule. We then directly compute the force using the worm-like-chain model, 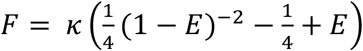

### Condensate nucleation probability analysis

To calculate the probability of the formation of a DNA-protein condensate as a function of end-to-end distance, we localized the peaks of the FoxA1 condensates from time-averaged projections of FoxA1 and DNA. We then extracted 0.9 *μ*m x 0.5 *μ*m windows centered around the localized FoxA1 peaks of both the FoxA1 and DNA signals—with the window’s long axis going with the strand and the short axis as orthogonal to the strand. We then computed the zero-lag normalized cross-correlation coefficient as follows:

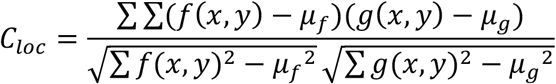

where *f*(*x, y*) is the DNA, *g*(*x, y*) is FoxA1, *μ*_*f*_ is the mean of the DNA image, and *μ*_*g*_is the mean of the FoxA1 image. This generates values from -1 to 1. For FoxA1- mediated DNA condensation, the values for particular condensates are close to 1. When FoxA1 fails to condense DNA, owing to the morphology of the underlying DNA strand and the small number of pixels, we obtain values that range from -1 to roughly 0.5. To obtain a value for *P*_*cond*_ as a function of end-to-end distance, we selected a threshold of 0.75—*C*_*loc*_values above the threshold are considered as “condensed” and values below would be considered “non-condensed”. We binned the *C*_*loc*_ data in 2-*μ*m increments as a function of end-to-end distance, and calculated *P*_*cond*_ by taking the number of “condensed” condensates and dividing it by the total number of condensates within the bin. The confidence intervals for *P*_*cond*_ in each respective bin are computed by computing the 95% confidence interval of a beta-distribution, which represents the probability distribution for a Bernoulli process that takes into account the total number of successes with respect to the total number of attempts.

### Parameter fitting of the thermodynamic description and confidence intervals

To fit *α*, we used a linear fit of the condensate volumes for individual strands as a function of *L*_*d*_. The confidence intervals are the 95 per cent CI generated from directly fitting the points. To fit the surface tension *γ* and condensation free energy per volume *ν*, we minimized the error of the average 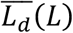, and *P*_*cond*_ (*L*) with respect to the data to optimize the parameter values. We used the normalized Boltzmann distribution 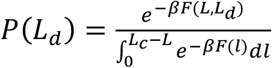 to calculate 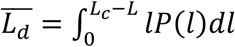. To compute *P*_*cond*_ (*L*), we localized the position of the local maximum in the free energy, 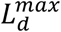, for a given L and then computed the probability to “not” nucleate a droplet from the Boltzmann distribution 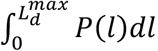,, which gives 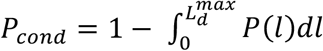. To minimize the error, we binned the data in 2-*μ*m-width bins. For each “binned” mean for both condensed DNA and condensation probability, we computed the squared residual of the mean value with respect to the theoretical expression. For residuals calculated from 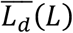,, we normalized each residual by the squared standard error of the mean, and then summed the normalized residuals to obtain the error. For residuals calculated from *P*_*nuc*_(*L*), we normalized each residual by the variance of the beta distribution, 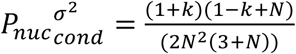, and then summed the normalized residuals to obtain the error. For the global error, we simply added the error from both deviations in 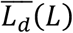 and *P*_*cond*_ (*L*). We then iterated through a range of values for (*γ, ν*) and computed the total error associated with each set of parameter values, exponentiated the negative values of the total error matrix, and computed the largest combined value to select the parameter values. To calculate the parameters’ confidence intervals, we obtained one-dimensional profiles of the integrated exponentiated total error for *ν* as a function of *γ* and *γ* as a function of *ν*. The peaks of these profiles represented the values that we selected for our best-fit parameters. We assumed that these profiles represented probability distributions for parameter selection, and then calculated the left and right bounds where the area under the curve between these bounds *r*epresented 95 per cent of the area. These left and right bounds represent the lower and upper values of our confidence intervals. To compute the 95 per cent confidence intervals for the force for each respective end-to-end distance value, we scanned through (*γ, ν*) parameter space and computed the value of *L*_*d*_ for each set of parameters. We then plotted these values against the probability that these parameter values were the “true” values—simply the probability from the exponentiated error matrix. Integrating the points under the Probability vs. *L*_*d*_ curve and dividing this by the total area under this curve, we generated a probability distribution function from which we could compute the 95% confidence intervals for *L*_*d*_. Because the force was constant, to compute the confidence intervals for the force, we calculated the force using the worm-like chain model using corresponding *L*_*d*_ values for an end-to-end distance that retained FoxA1-mediated DNA condensation. To compute the confidence intervals for *L*_*crit*_, we scanned through (*γ, ν*) parameter space and computed *L*_*crit*_ for each set of parameters. We then plotted *L*_*crit*_ values with the corresponding values from the probability that these parameter values were true (again, the exponentiated error matrix). Integrating the points under the Probability vs *L*_*crit*_ curve and dividing this by the total area under this curve, we generated a probability distribution function from which we could compute the 95% confidence intervals for *L*_*crit*_.

## Data availability statement

Source data files are made available for this paper. Data generated and analysed supporting the findings of this manuscript will be made available upon reasonable request.

## Code availability statement

Code generated supporting the findings of this manuscript will be made available upon reasonable request.

**Extended Data Figure 1:**
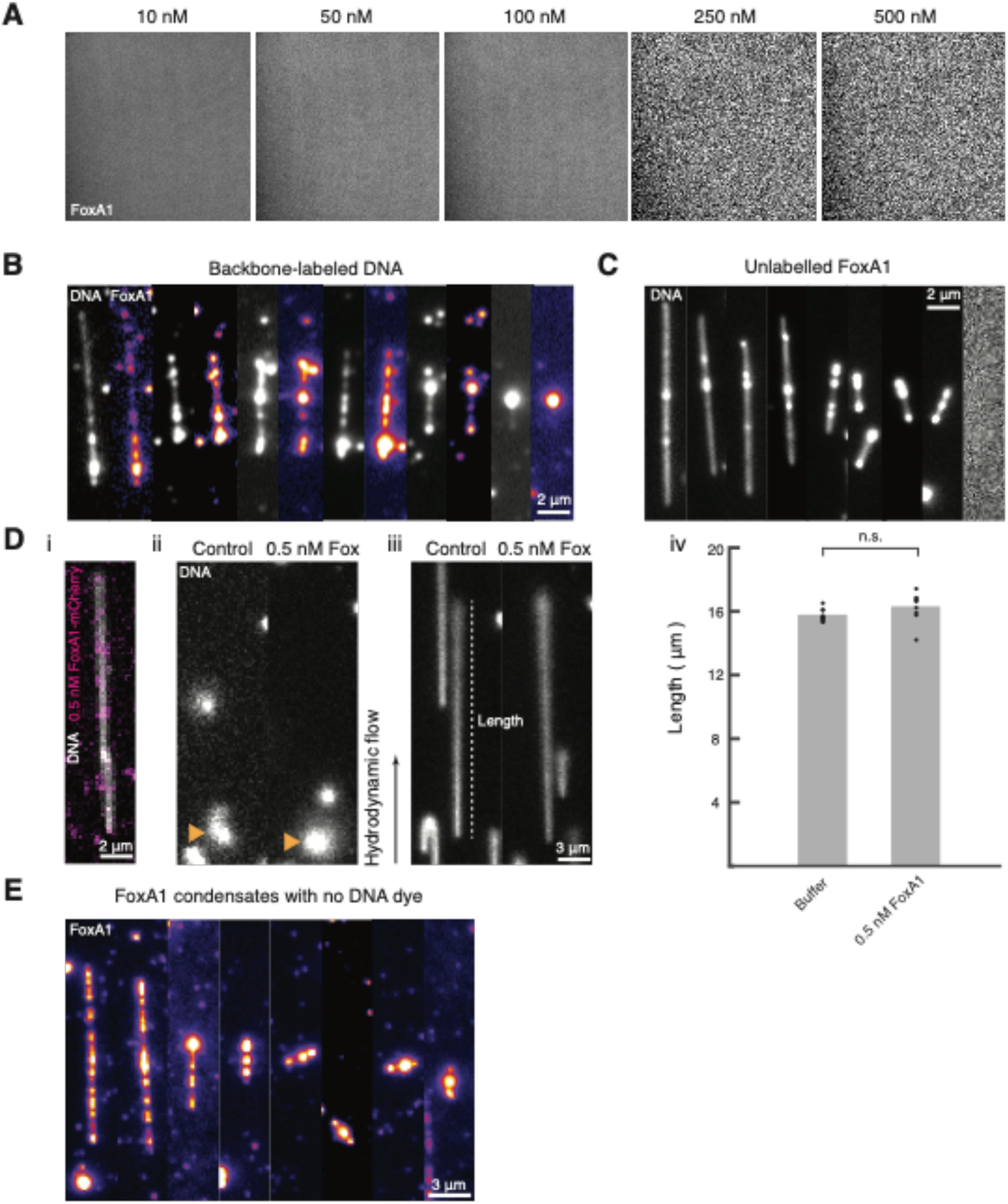
Experimental controls for FoxA1-mediated DNA condensation. (A) Representative fluorescent images of FoxA1-mCherry in buffer (20 mM HEPES, pH=7.8, 50 mM KCl, 2 mM DTT, 5% glycerol, 100 μg/ml BSA) at different concentrations, 10-500 nM, in the absence of DNA reveals that FoxA1 does not form condensates in bulk at these concentrations. Using spinning disk microscopy and a 60x objective, we acquired images 70 *μ*m x 70 *μ*m in size with an exposure time of 250 msec and a time stamp of 500 msec to generate movies 30 seconds in duration. For all measured concentrations we generated n=3 movies and did not observe any FoxA1 condensation. (B) FoxA1-mCherry condenses λ- phage DNA molecules with Cy5 dye covalently attached to the phosphate backbone of DNA (Label-IT Nucleic Acid Labeling Kit, Cy5, Mirus). (C) Unlabeled FoxA1 condenses DNA (visualized with 10 nM Sytox Green). The rightmost panel is a representative image of the mCherry 561 nM imaging channel, revealing that the FoxA1 molecule does not have a mCherry fluorophore. (D) Sparse labeling of FoxA1 (0.5 nM) does not influence the persistence length and contour length of λ- phage DNA, as determined by hydrodynamic stretching (see Methods). (i) FoxA1 (purple) is sparsely bound to DNA (in grey), visualized with 10 nM Sytox Green. (ii) Snapshots of unstretched DNA molecules bound at only one end to the coverslip before hydrodynamic stretching in both control and 0.5 nM FoxA1 conditions. The yellow arrows point to the DNA molecules. (iii) Snapshots of stretched DNA molecules bound at one end to the coverslip during hydrodynamic stretching in both control and 0.5 nM FoxA1 conditions. (iv) Quantification of stretched DNA lengths in both control (n=10) and 0.5 nM FoxA1 (n=9) conditions reveals that there is no significant difference in the length under hydrodynamic stretching (unpaired t-test, p=0.11). (E) FoxA1 condensates imaged in the absence of DNA dye are consistent in size with that of FoxA1 condensates formed in the presence of DNA dye.

**Extended Data Figure 2:**
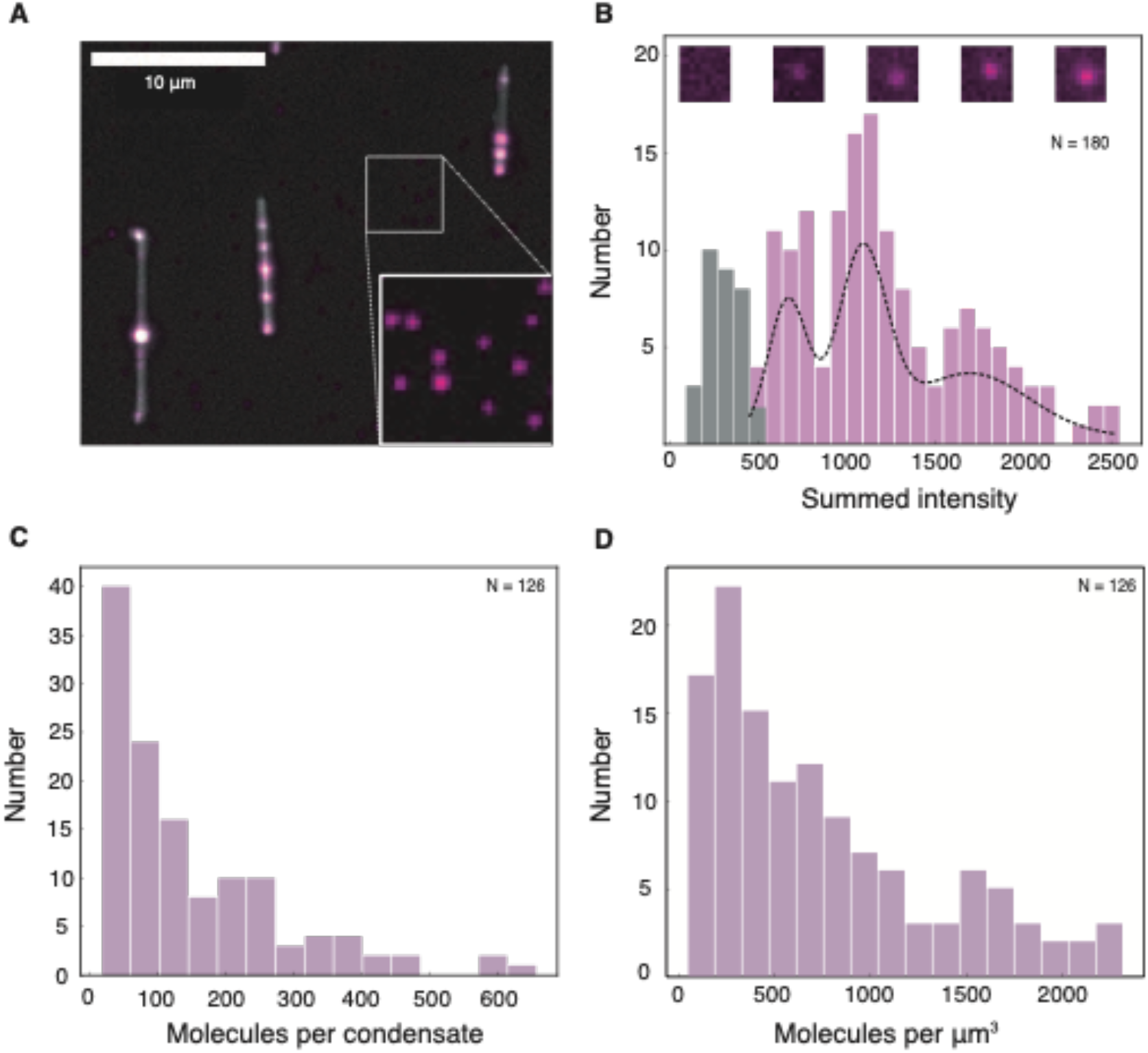
Counting FoxA1 molecules in condensates. (A) Representative image of three DNA strands with FoxA1 condensates. The inset shows an area of the PEGylated glass slide void of DNA. Increased contrast reveals the presence of individual spots of FoxA1 non-specifically bound to the coverslip. (B) Histogram of integrated intensities of these DNA-independent FoxA1 to calibrate the amount of fluorescence per molecule. The grey bars represent the integrated background intensity of areas where no FoxA1 signal could be detected (maximum at 289 a.u.). Pink bars represent the integrated intensity of individual spots of DNA-independent FoxA1 signal. Black dotted line is a multi-Gaussian fit to the pink histogram, indicating consecutive peaks in the histogram at intensities of 683, 1096 and 1706 (a.u.), suggesting an integrated intensity of 400 a.u. per FoxA1 molecule. Representative images (10×10 pixels) of background (left) and individual DNA-independent FoxA1 spots used in this analysis are placed above the histogram according to their integrated signal intensity. (C) Histogram of the number of FoxA1 molecules in FoxA1 condensates on DNA, calculated based on an integrated intensity of 400 a.u. per FoxA1 molecule, determined in (B). The mean number of molecules is 150 per condensate. (D) Histogram of the density of FoxA1 molecules in the FoxA1-DNA condensates analyzed in (C). The mean value is 750 molecules per μm^3^. These estimates represent lower bounds as previous studies have demonstrated that fluorescent-based methods for estimating the number of molecules neglect effects such as quenching and can underestimate the number of molecules by as much as 50 fold^27^.

**Extended Data Figure 3:**
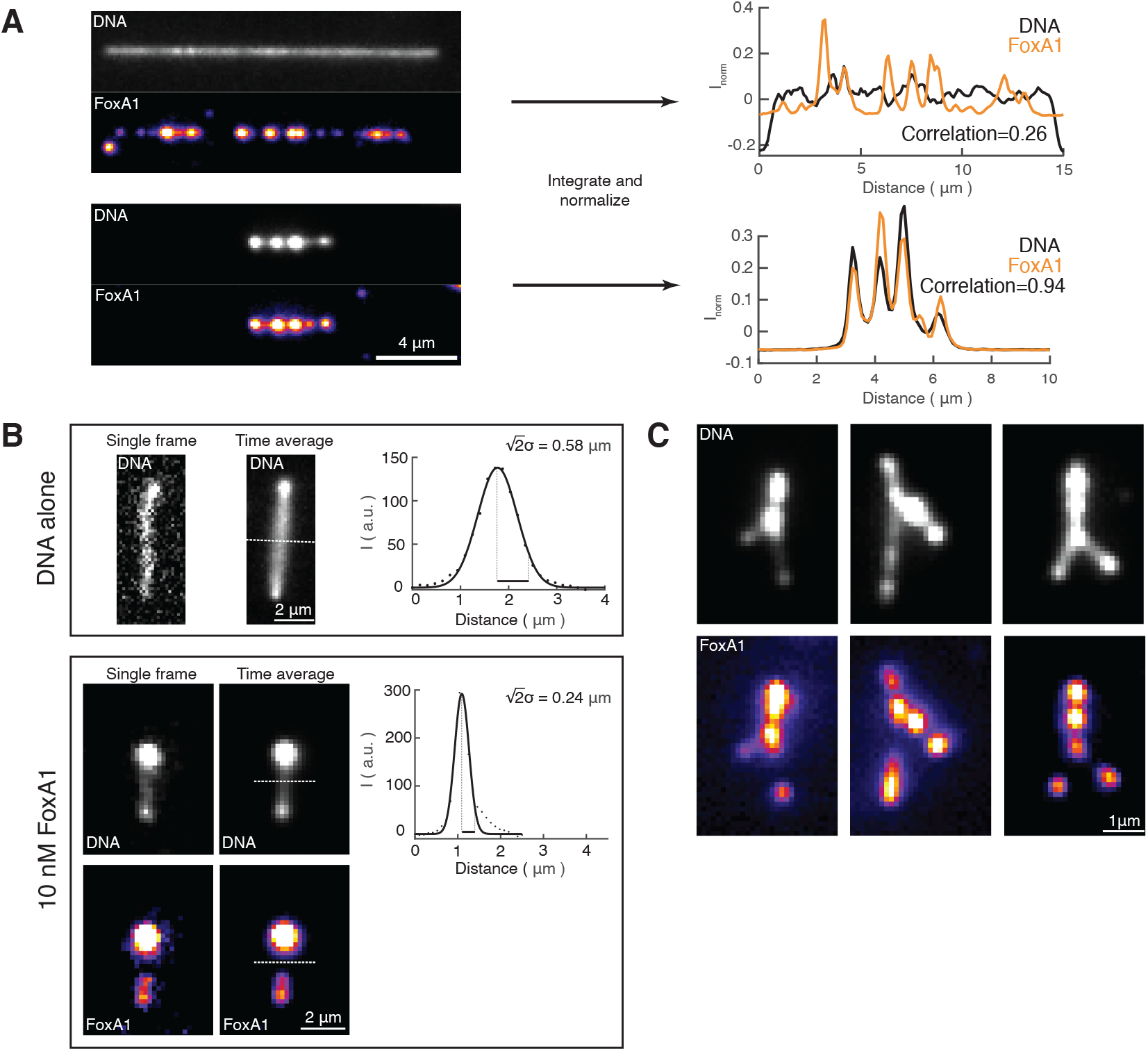
Quantification of FoxA1-mediated DNA condensation. (A) Global cross-correlation between FoxA1 and DNA reveals FoxA1-mediatd DNA condensation. Left, representative fluorescent time-averaged projections of DNA and FoxA1 at two different end-to-end distances. Integrating both the DNA and FoxA1 signals along the axis orthogonal to the long axis of the strand gave rise to line profiles, which we normalized, and then plotted as a function of distance (DNA in black and FoxA1 in orange). We then computed the zero-lag cross-correlation coefficient defined as “Correlation” (see Methods). (B) DNA envelope width measure measures the transverse fluctuation of non-condensed DNA. Top box: DNA alone condition. Bottom box: DNA+FoxA1 condition. For both conditions, we display representative fluorescent images of single frames and time-averaged projections of the DNA and FoxA1 signals. The white dashed line represents the maximum width of the DNA signal along the orthogonal axis of the non-condensed DNA. The black dots in the profile represent the background-subtracted points from the white dashed line, and the black line represents a Gaussian fit. The DNA envelope width was defined as 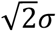,, where *σ* is the standard deviation of the Gaussian fit. (C) Three representative examples of FoxA1-mediated zipping. These images are time-averaged projections of both FoxA1 and DNA.

**Extended Data Figure 4:**
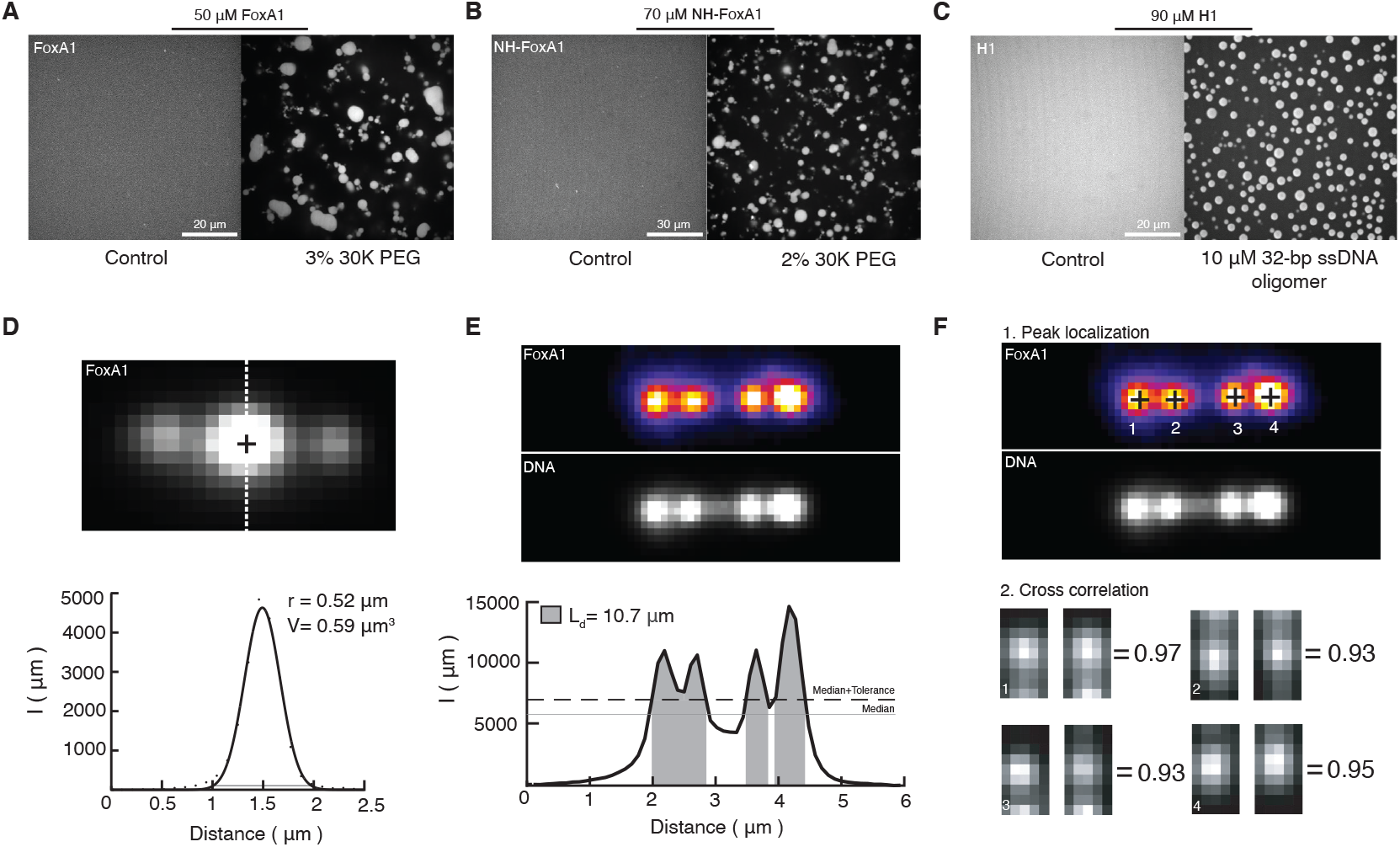
Bulk biomolecular condensate formation and quantification of condensate volume, condensed DNA length, and condensation probability. (A) Three per cent 30K PEG triggers FoxA1 condensate formation in bulk at 50 μM in storage buffer: 20 mM HEPES (pH=6.5), 100 mM KCl, 1 mM MgCl2, 3 mM DTT, and 2 M Urea. (B) Two per cent 30K PEG triggers NH-FoxA1 condensate formation in bulk at 70 μM in storage buffer. (C) The addition of 10 μM 32-BP ssDNA oligomers nucleated droplets of H1 in bulk at 90 μM that exhibited features of liquid-like droplets consistent with literature^28,29^. These data demonstrate that H1-DNA form liquid-like condensates, which could be driven via transient cross-linking of H1 and DNA or H1-H1 interactions. Both mechanisms are accounted for in our free energy description. (D) Condensate volume quantification of a representative time-averaged projection of a FoxA1- DNA condensate, where the black cross is the condensate peak location and the white dashed line is the intersecting profile to measure the volume. Lower panel: the black dots are the profile’s background-subtracted values and the solid black line is a Gaussian fit. The gray line represents the threshold value computed from the gradient of the Gaussian function that defines the edges of the condensate (see Methods). (E) Condensed DNA length quantification of a representative time-averaged projection of FoxA1 and DNA. Below: the integrated one-dimensional DNA profile is defined into condensed versus non-condensed regions using the median of the profile’s median (gray) plus a tolerance (black dashed). (F) Local correlation quantification of a representative time-averaged projection of FoxA1 and DNA. The condensates were localized (black crosses) and then 0.9 μm x 0.5 μm boxes centered around these peaks were cropped. The correlations between the cropped regions of FoxA1 (left) and DNA (right) were then computed.

**Extended Data Figure 5:**
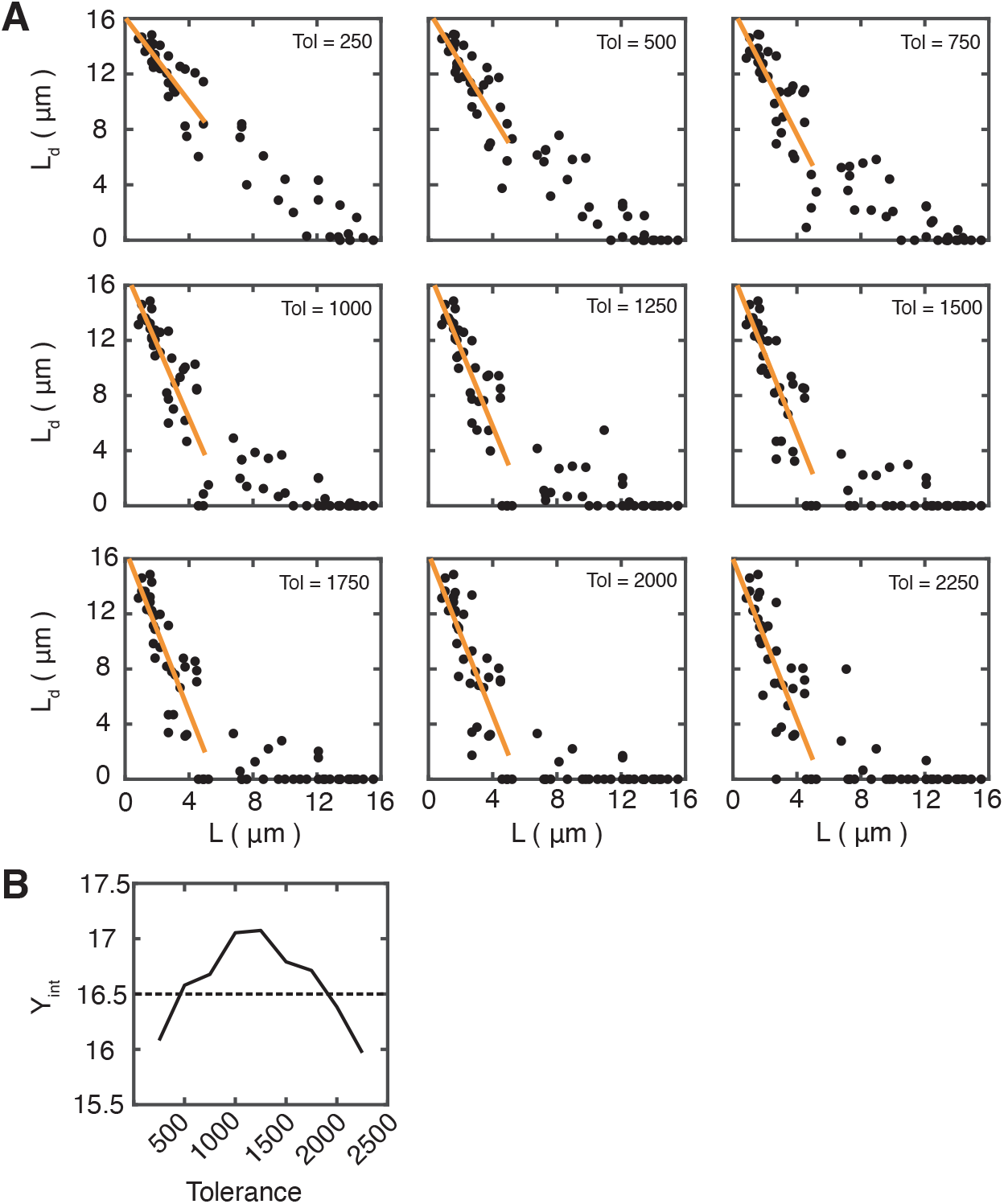
Tolerance value calculation. Quantification of the condensed DNA length as a function of end-to-end distance for a range of tolerance values. Condensed DNA length is computed by defining regions of condensed versus non-condensed DNA using a threshold composed of the signal’s median value plus a tolerance. (A) Condensed DNA length is plotted as a function of end-to-end distance L for tolerance values from 250 to 2250 where the black dots represent the condensed DNA length for individual strands and the orange curve represent linear fits to these points for end-to-end distance below 5 μm. (B) Y intercept of the fitted linear curves. A tolerance=500 was selected as the y intercept was equal to the contour length of λ-phage DNA (16.5 μm) and gave FoxA1-DNA condensate formation up to approximately 10 μm, consistent with experimental observations (see Methods).

**Extended Data Figure 6:**
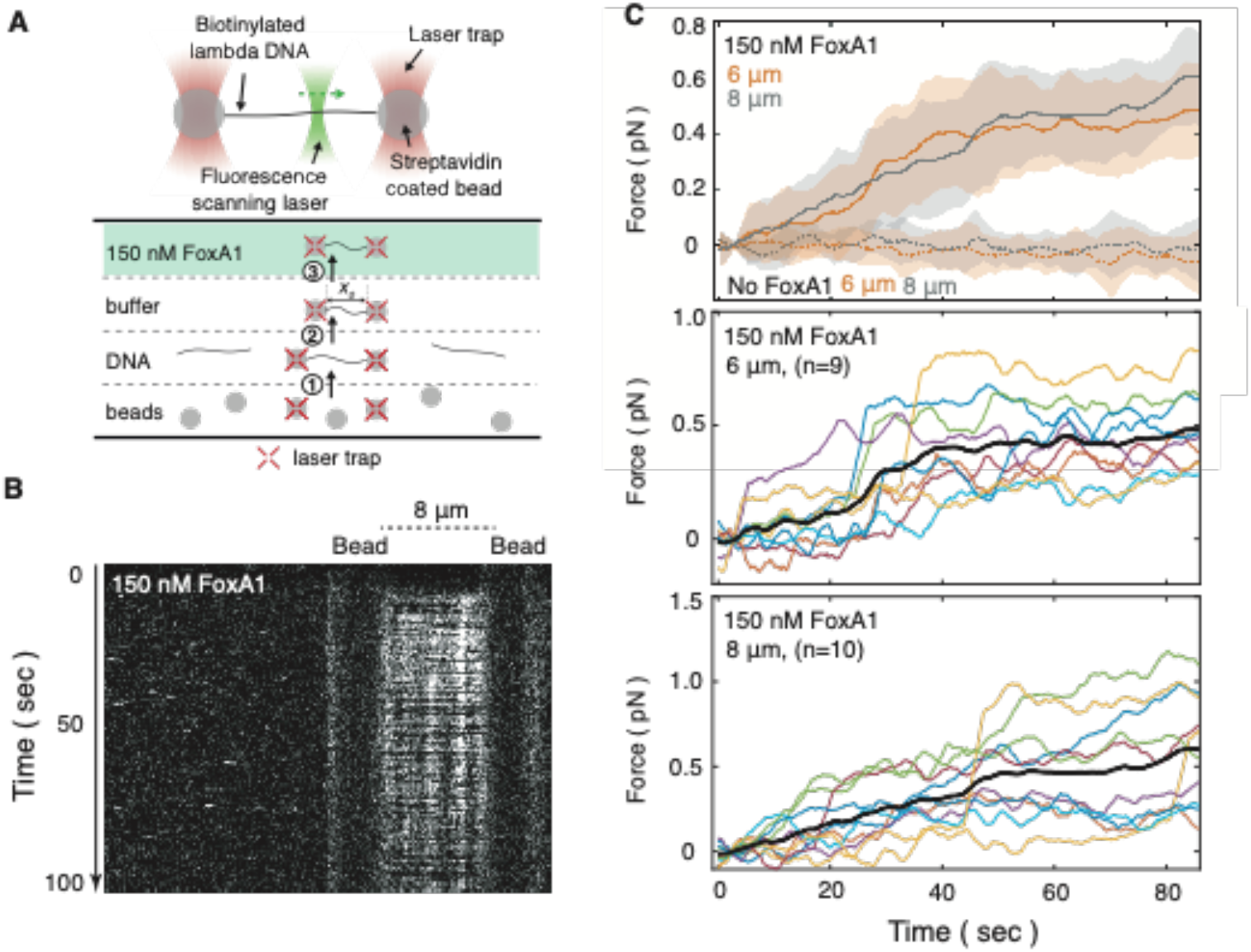
Optical tweezer measurements reveal that FoxA1 generates forces on the order of 0.4-0.6 pN. (A) Schematic outlining optical tweezer experimental design (see Methods). (B) Representative kymograph reveals that FoxA1 condensates co-localize with a single molecule of λ-phage DNA trapped between two beads at an end-to-end distance of 8 μm. (C) Force trajectories for single DNA molecules reveal forces on the order of 0.4-0.6 pN when in FoxA1-containing buffer. (Top panel) This panel displays the mean ± STD of force trajectories for each condition (n=9 for +FoxA1 with L=6 μm, n=10 for +FoxA1 with L=8 μm, n=10 for control with L=6 μm, and n=13 for control with L=8 μm.). This average force is slightly higher than what we measured in Fig. 3F using fluorescence, though a comparison of the relative errors reveals that both measurements give rise to comparable forces close to their respective detection limits and within the error bars. Additionally, the optical tweezer measurements were performed at a higher FoxA1 concentration—this was due to the large amount of tubing from the entry port to the flowcell in the custom-built Lumicks system, representing a considerable amount of surface for the protein to non-specifically bind to. We found that 150 nM FoxA1 was necessary to elicit a force response and to observe FoxA1 condensate formation on DNA. We conducted these measurements in the presence of 150 nM FoxA1 in Cirillo buffer 20 mM HEPES, pH=7.8, 50 mM KCl, 3 mM DTT, 5% glycerol, 100 μg/ml BSA (solid lines) and in the presence of Cirillo buffer only (hatched lines) at end-to-end distances of L=6 (orange) or 8 μm (grey). Individual force trajectories for λ-phage DNA in the presence of buffer containing 150 nM FoxA1 with an initial end-to-end distance of 6 μm (middle panel) and 8 μm (bottom panel) reveal jumps in force, consistent with a first-order phase transition. These trajectories are re-plotted for clarity in Extended Data Fig. 7.

**Extended Data Figure 7:**
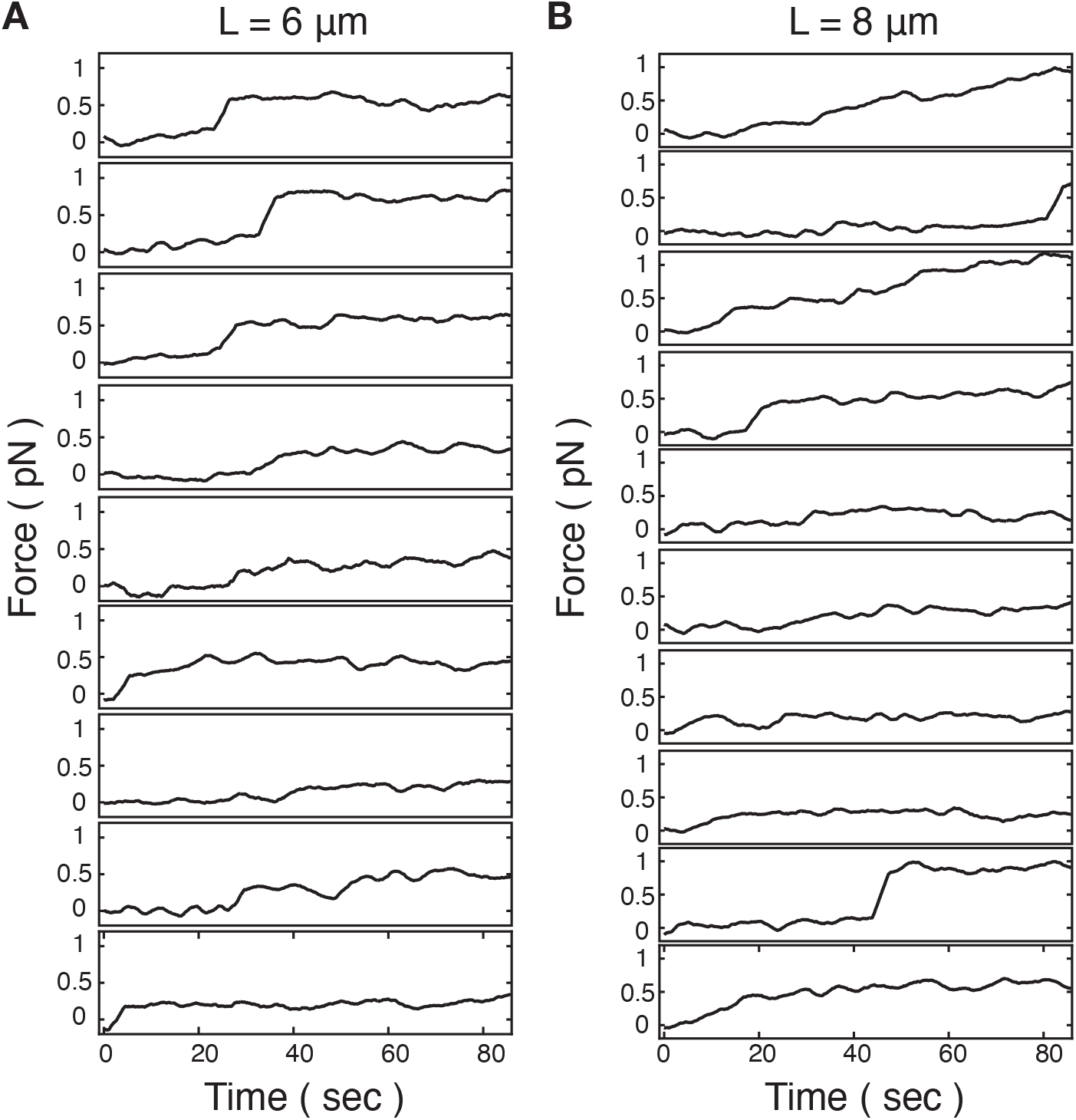
Individual temporal optical tweezer force measurements. Temporal force measurements from optical tweezers with an initial end-to-end distance of 6 μm (n=9 strands) (A) and 8 μm (n=10 strands) (B) in the presence of 150 nM FoxA1. These data are the same as in Extended Data Fig. 6c, and are re-plotted individually for clarity.

**Extended Data Figure 8:**
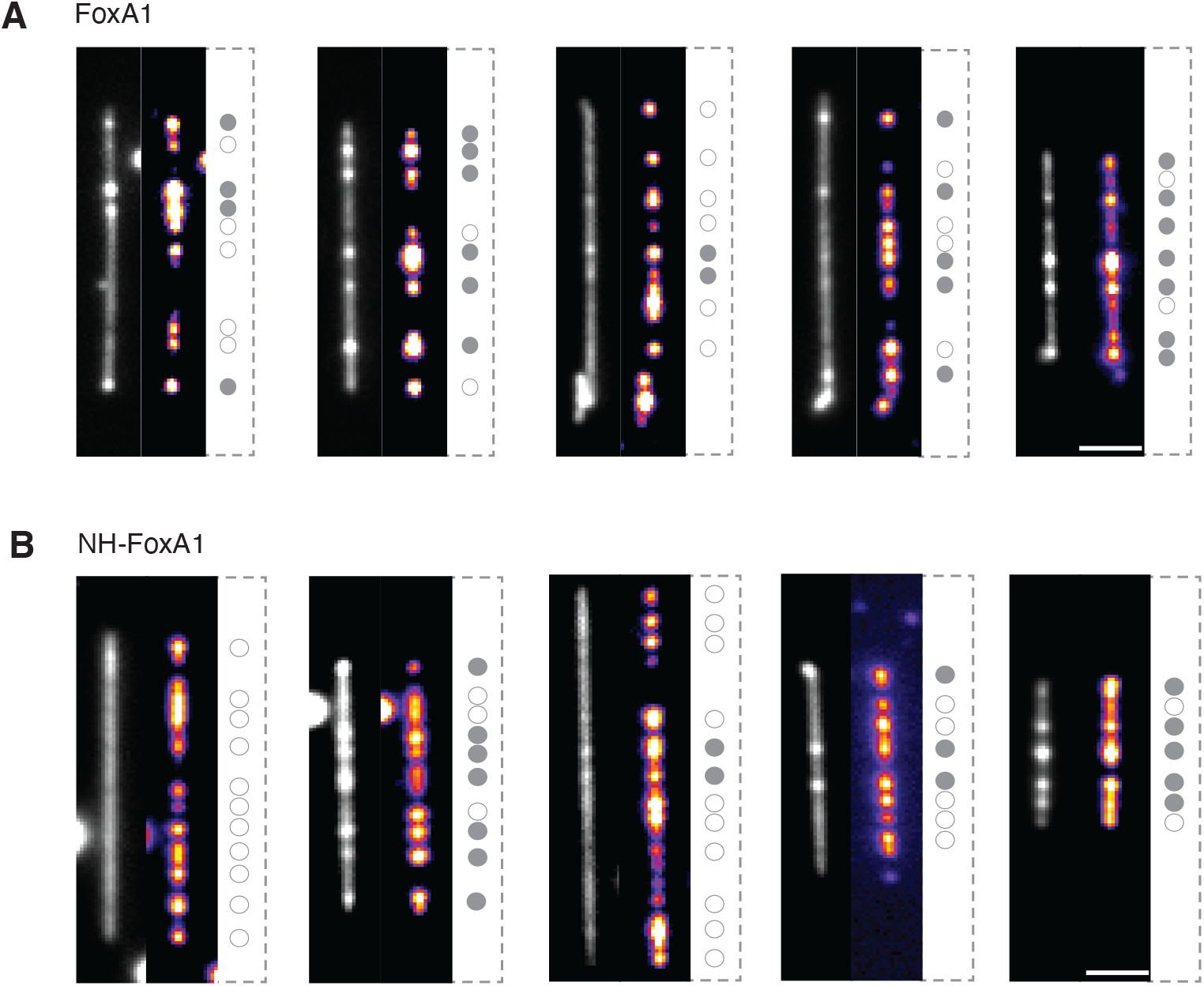
Bistability of FoxA1-mediated DNA condensation. (A) Representative time-averaged projections of DNA and FoxA1 signals show that FoxA1 condenses DNA in an all-or-nothing manner. On the right side of each pair of images, we localized the FoxA1 condensates and showed whether FoxA1 condenses DNA (filled-in gray circle) or not (open circle). Interestingly, there is a mixed population, revealing the bistable nature of the condensation process. (B) Representative images of condensation bistability for the sequence-specific DNA- binding mutant, NH-FoxA1. Scale bars = 2 μm.

**Extended Data Figure 9:**
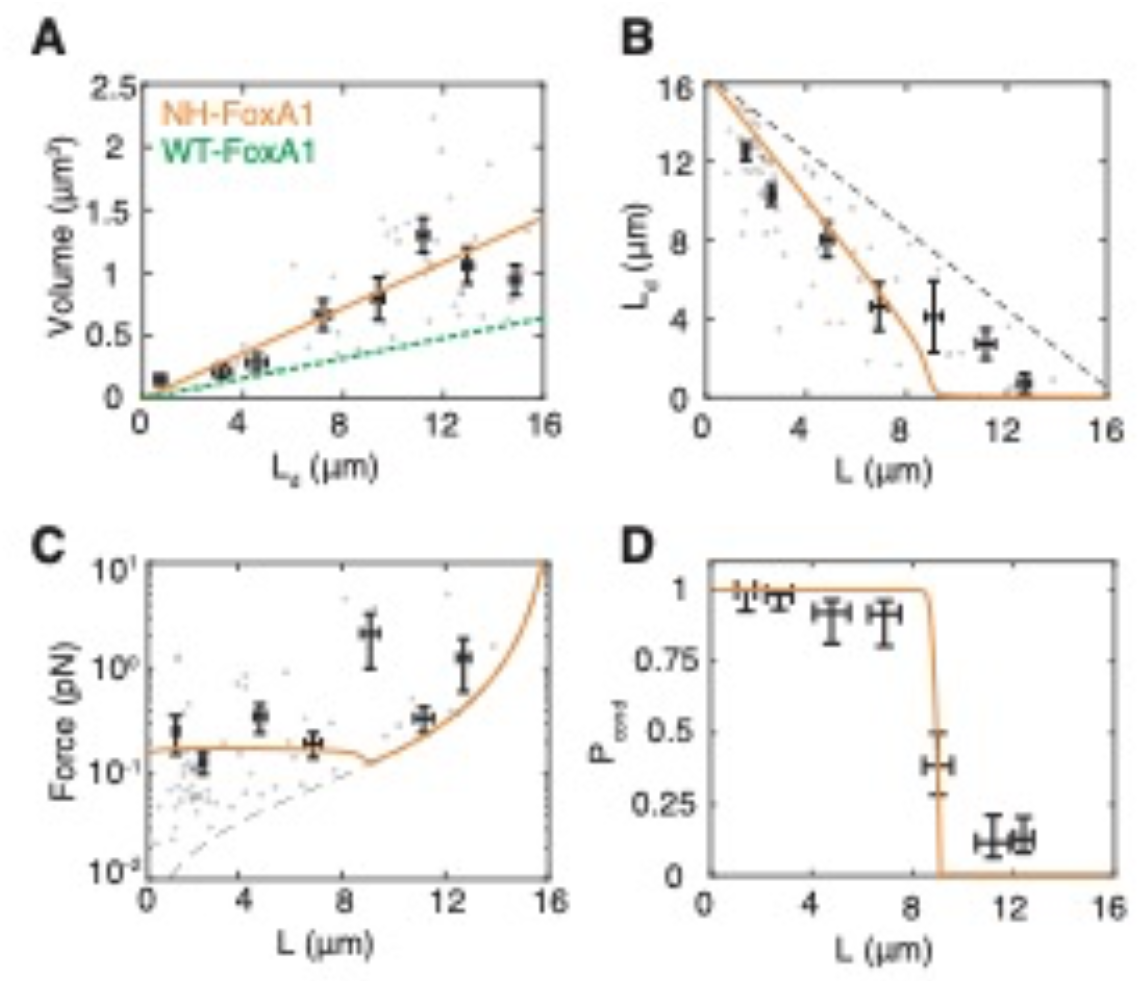
Quantification of NH-FoxA1-mediated DNA condensation. (A) Condensate volume as a function of condensed DNA length (L_d_). The grey dots represent individual strands (n=47) and the data is binned every 2 μm (mean ± SEM). The individual data are points are fit with a linear curve with a slope of 0.09 μm^2^ given in orange. The green dashed line is the WT-FoxA1 fit (slope=0.04 μm^2^). (B) Condensed DNA length as a function of end-to-end distance. The black dots represent individual strands (n=70) and the data is binned every 2 μm (mean ± SEM). The orange curve is the expression computed from the theoretical description with parameter values determined through error minimization (see Methods). The black hatched line represents the DNA’s contour length (16.5 μm) minus the end-to-end distance. (C) The force that the condensate exerts on the non-condensed DNA as a function of end-to-end distance. The grey dots represent individual strands (n=68) and the data is binned every 2 μm (mean ± SEM). The orange curve is the expression computed from the theoretical expression of L_d_ versus L from panel B for the force. NH-FoxA1 generates forces at roughly 0.17 pN. The dashed black line represents the force exerted on the non-condensed strand when L_d_=0. (D) Probability for NH-FoxA1 to form a DNA-FoxA1 condensate reveals a sharp transition at a critical end-to-end distance. Local correlations of individual FoxA1 condensates with DNA (Extended Data Fig. 4c) are calculated, binned into 2-μm-width bins, and *P*_*cond*_ is calculated (see Methods). There are a total number of n=361 condensates used for this analysis. The dashed lines represent the *P*_*cond*_ value as computed within the bin with ± SD for the strand’s end-to-end distance. The confidence intervals for *P*_*cond*_ are computed by computing the 95% confidence interval of a beta-distribution (see Methods). The orange curve represents *P*_*cond*_ computed from the theoretical description with parameter values determined through error minimization.

**Extended Data Figure 10:**
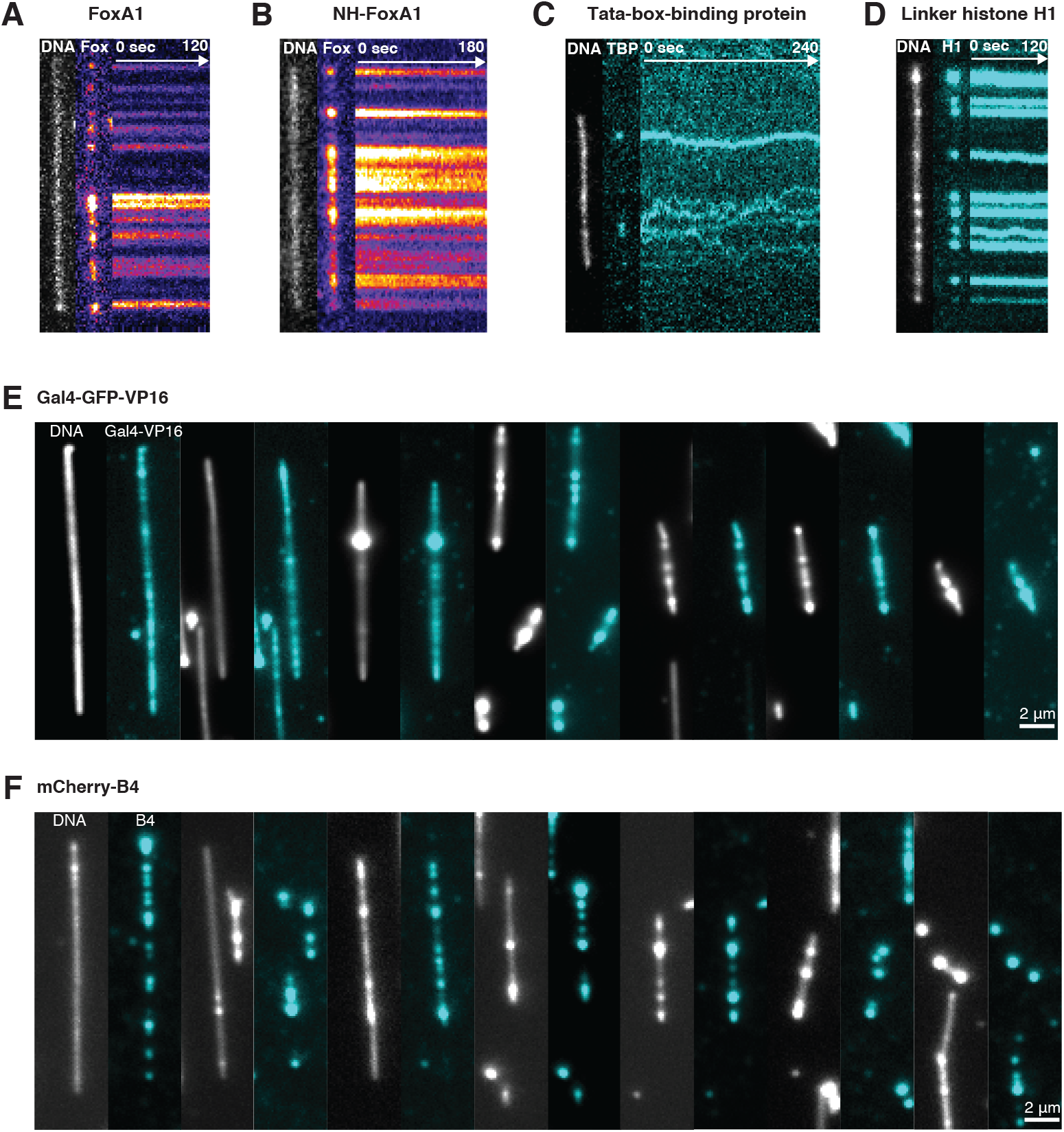
Dynamics of DNA-binding proteins. (A) Representative images of FoxA1 condensates on DNA. The kymograph reveals FoxA1 condensates do not move on DNA. (B) NH-FoxA1 condensates remain stable on DNA and do not move. (C) TBP condensates exhibit diffusive-like behavior on DNA. (D) Similar to FoxA1 condensation, H1 condensates do not exhibit diffusive-like behavior on DNA. (E) Representative images of Gal4-GFP- VP16-mediated DNA condensation. DNA was imaged with 10 nM Sytox Orange. (F) Representative images of mCherry-B4-mediated DNA condensation. DNA was imaged with 10 nM Sytox Green.

## Supplementary Information

### 1 Thermodynamic description of DNA-protein condensation

We consider the free energy associated with nucleating a condensate that contains DNA and FoxA1. The free energy of this process contains volume and surface contributions of the DNA- protein condensate as well as the free energy of the DNA polymer outside the condensate,

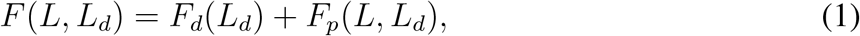

where *L* is the end-to-end distance of the DNA, *L*_*d*_ is the length of condensed DNA, *F*_*d*_ is the free energy of the condensate, and *F*_*p*_ is the free energy of the DNA polymer outside the condensate. Assuming that the DNA co-condenses with the protein to form a dense condensed phase with defined volume fraction of DNA, the droplet volume and the length of condensed DNA are linearly related, *V* = *αL*_*d*_, or 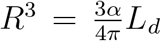, where 1*/α* describes the DNA packing density given as DNA length per condensate volume. We can then obtain the condensate free energy of nucleating a condensate as a function of *L*_*d*_ and end-to-end distance *L* as

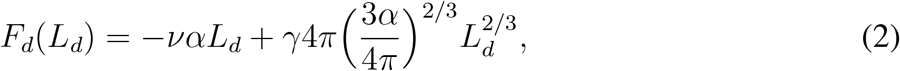

where *ν* is the condensation free energy per volume, and *γ* is the surface tension of the condensate. The free energy of the polymer *F*_*p*_(*L, L*_*d*_) is related to the external force applied to pin the free DNA polymer and its associated chemical potential by

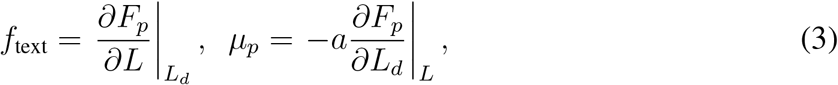

where *a* is the length of a base pair. The force-extension relation for *λ*-phage DNA has been extensively studied previously, and here we use the phenomenological force-extension curve of the worm-like-chain model for *λ*-phage DNA (14) with contour length, *L*_*c*_ (for *λ*-phage DNA *L*_*c*_ = 16.5 *µ*m). If a length *L*_*d*_ of the DNA is condensed, the extension of the non-condensed strand is 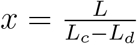. The force on the strand then can be expressed as

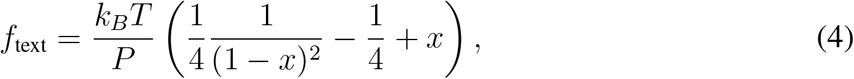

where *k*_*B*_ is the Boltzmann constant, *T* is the temperature, and *P* is the persistence length of DNA. For what follows we define 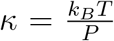. From this expression of the force and its relation to the free energy of the DNA polymer (equation 3), we can obtain the free energy of the DNA polymer outside the condensate as 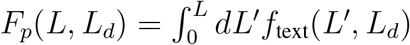, leading to

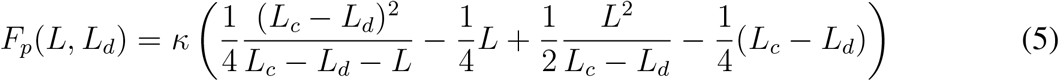

The total free energy associated with nucleating a FoxA1-DNA condensate on a DNA strand reads:

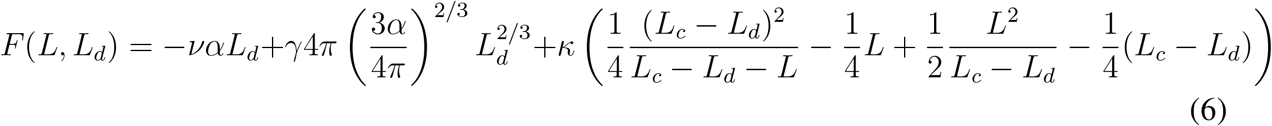

The equilibrium between condensate and polymer is given by 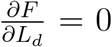,, which is equivalent to equilibrating the chemical potentials of the condensate and free polymer,

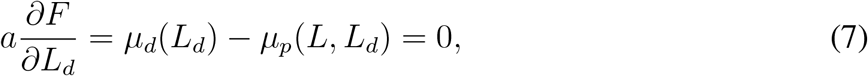

with

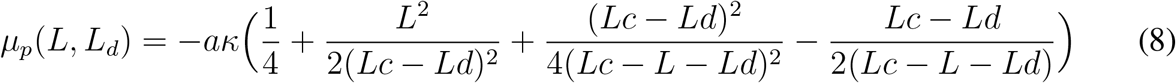

Using the expression for the total free energy, we can vary the length *L*_*d*_ of condensed polymer and obtain profiles for the free energy as a function of *L*_*d*_, which depend on the end-to-end distance *L* (see Fig. 3b). For *L* values close to 0—where the strand is not under tension—we observe that there is a minimum of *F* for *L*_*d*_ close to *L*_*c*_. This means that, at this end-to-end distance, FoxA1 has mediated the generation of a FoxA1-DNA condensate using almost all of the DNA in the strand. As *L* increases, however, the local minimum shifts to lower values of *L*_*d*_ and ultimately *F* at the minimum becomes higher than the free energy without condensate *F* (*L*_*d*_ = 0), giving rise to a branch of metastable states. For even higher *L* values, the metastable state disappears and the global minimum is at *L*_*d*_ = 0 (Fig. 3b). This sharp transition corresponds to a first-order phase transition. Simple scaling arguments are useful to generate intuition for the conditions necessary for condensate formation, and for the condensate to pull DNA. Briefly, there are three energy scales associated with this problem: the energy associated to create a droplet, which is *ναL*; the surface energy of scale 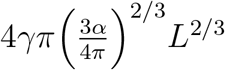; and lastly the energy scale associated to the non-condensed polymer 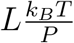. First, to create a droplet,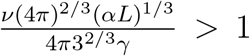. Once condensation is favorable, in order for the droplet to pull DNA, 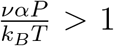. Notably, fitting the parameter values (see Methods) demonstrated that, at low *L*, the free energy gained by the system is on order of 1-2 *k*_*B*_*T*, implying that stochasticity is relevant for the condensation process. To account for the inherent stochastic nature of the condensation, we compute the probability of nucleating a DNA-protein condensate of size *L*_*d*_ using Boltzmann distributions from the corresponding energy profiles,

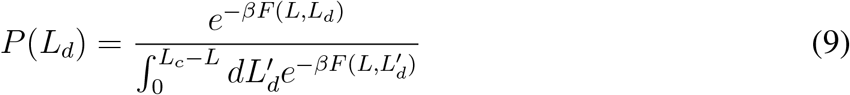

where 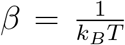. To determine the relationship between *L*_*d*_ and *L*, we compute the mean *L*_*d*_ value of these Boltzmann distributions: 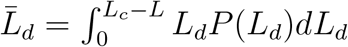 which then allows us also to calculate the magnitude of the condensation forces using the worm-like chain model given in Eq. (4).

### 2 Supplementary Information Figures

**Figure 1:**
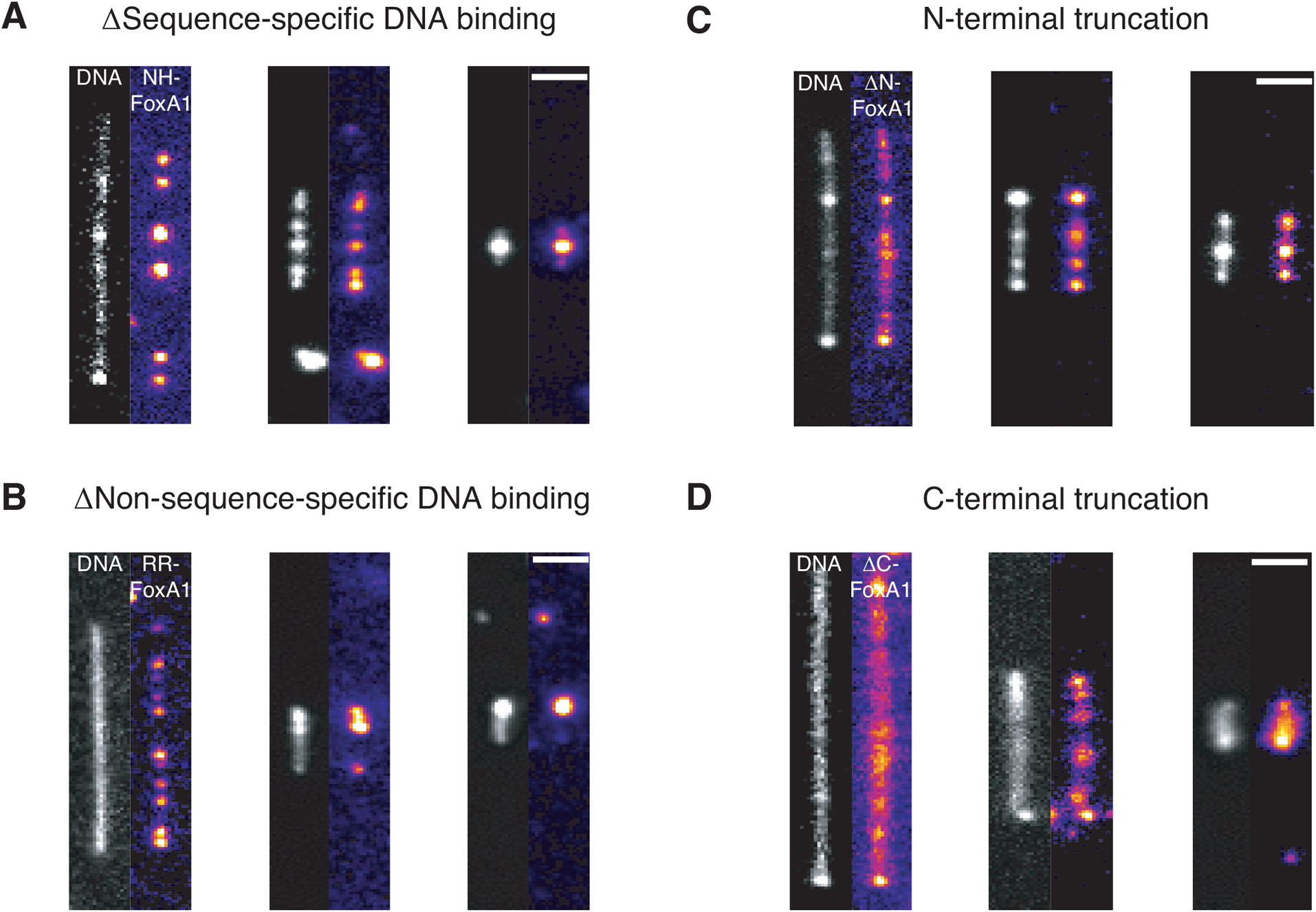
Representative images of sequence-specific-binding NH-FoxA1 mutant (A) non-sequence-specific-binding RR-FoxA1 mutant (B) N-terminal FoxA1 truncation (C) and C-terminal FoxA1 truncation (D). The scale bars are 2 *µ*m. DNA is imaged with 10 nM Sytox Green. Note that the contour length of each DNA molecules is constant (16.5 *µ*m) but the end-to-end distance is different.

**Figure 2:**
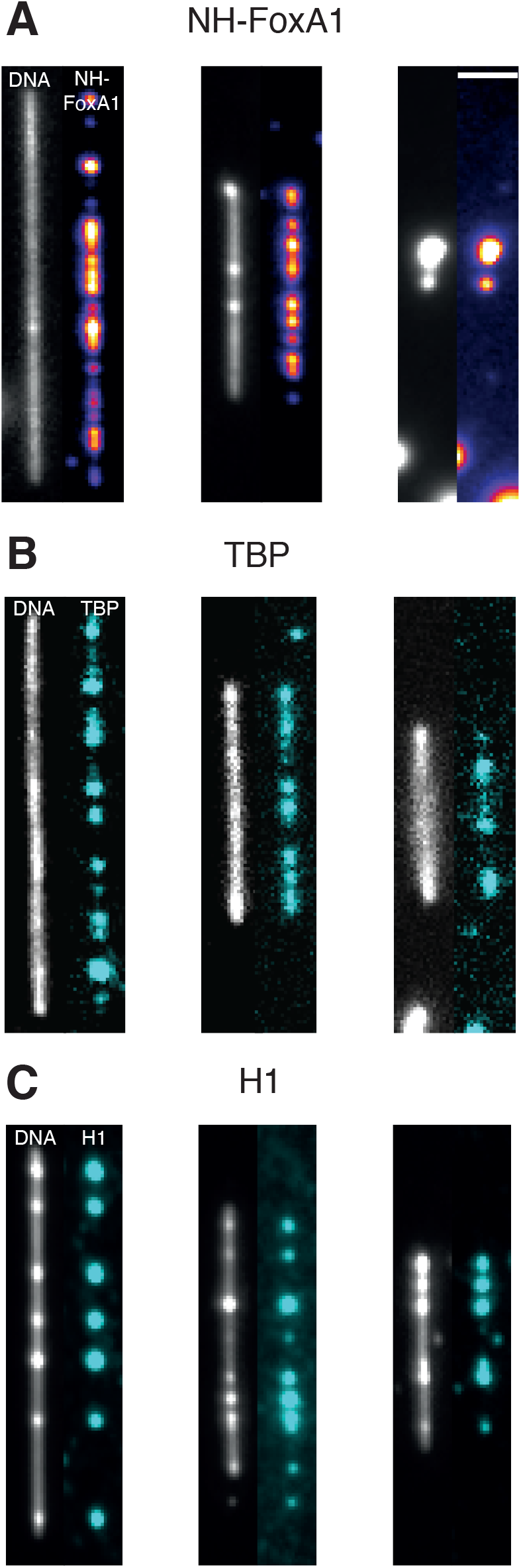
Representative images for NH-FoxA1 (A) Tata-box-binding protein (B) and somatic linker histone H1 (C). The images are time-averaged projections of movies for NH-FoxA1 and H1 but single images for TBP owing to TBP’s diffusivity. The scale bar is 2 *µ*m.

